# Does size matter? The relationship between predictive power of single-subject morphometric networks to spatial scale and edge weight

**DOI:** 10.1101/170381

**Authors:** Pradeep Reddy Raamana, Stephen C. Strother, for the Australian Imaging Biomarkers and Lifestyle flagship study of ageing, for The Alzheimer’s Disease Neuroimaging Initiative

**Author notes:** Data used in the preparation of this article was obtained from: 1) Alzheimer Disease Neuroimaging Initiative (ADNI) and 2) the Australian Imaging Biomarkers and Lifestyle flagship study of ageing (AIBL) funded by the Commonwealth Scientific and Industrial Research Organisation (CSIRO) which was made available at the ADNI database (www.loni.usc.edu/ADNI). The ADNI and AIBL researchers contributed data but did not participate in analysis or writing of this report.

## Abstract

Network-level analysis based on anatomical, pairwise similarities (e.g., cortical thickness) has been gaining increasing attention recently. However, there has not been a systematic study of the impact of spatial scale and edge definitions on predictive performance. In order to obtain a clear understanding of relative performance, there is a need for systematic comparison. In this study, we present a histogram-based approach to construct subject-wise weighted networks that enable a principled comparison across different methods of network analysis. We design several weighted networks based on three large publicly available datasets and perform a robust evaluation of their predictive power under four levels of separability. An interesting insight generated is that changes in nodal size (spatial scale) have no significant impact on predictive power among the three classification experiments and two disease cohorts studied, i.e., mild cognitive impairment and Alzheimer’s disease from ADNI, and Autism from the ABIDE dataset. We also release an open source python package called *graynet* to enable others to implement the novel network feature extraction algorithm, which is applicable to other modalities as well (due to its domain- and feature-agnostic nature) in diverse applications of connectivity research. In addition, the findings from the ADNI dataset are replicated in the AIBL dataset using an open source machine learning tool called *neuropredict*.

## Introduction

Network-level analyses have become one of the dominant techniques to process and analyze different neuroimaging modalities, including functional MRI (task- and resting-state fMRI), and diffusion MRI. One of the most routine network analyses performed is the extraction of individual connectivity matrices e.g. to characterize the structure and function of the brain, as well as to develop markers of dysfunction in various brain disorders. Owing to their broad applicability and success, similar approaches have been developed in the structural MRI (sMRI) also (Pradeep Reddy Raamana et al. 2015). Translation of such powerful techniques to the sMRI, and a systematic evaluation of their methodological robustness, would help assess clinical utility, esp. in the development of computer-aided diagnostic (CAD) techniques for deadly brain disorders like the Alzheimer’s disease (AD) (Alzheimer’s Association 2017).

Although there has been great progress in the last few decades in accurately characterizing AD as well as its progression (Weiner et al. 2017; 2015), its translation to improvement of clinical trials continues to be a great challenge (Cummings, Morstorf, and Zhong 2014). For any preventive or disease-modifying therapies to succeed, early prognosis is key. Towards this goal, diverse regional and network-level analyses of features derived from different neuroimaging modalities such as sMRI (Cuingnet et al. 2011; Bron et al. 2015; Duchesne et al. 2008; P R Raamana et al. 2014; Dyrba et al. 2015), positron emission tomography (PET) (Dukart et al. 2011; Herholz et al. 2002; Matthews et al. 2016) and resting-state fMRI (Hojjati, Ebrahimzadeh, and Khazaee 2017; Abraham et al. 2017) have been developed and are showing great promise in identifying differences between health and disease in the early stages, as well as establishing how they correlate with cognitive measures (Alexander-Bloch, Giedd, and Bullmore 2013; Tijms et al. 2013). Multimodal predictive modeling methods typically demonstrate higher prognostic accuracy (Sui et al. 2011; Arbabshirani et al. 2017) in many applications, owing to their training based on multiple sets of rich and complementary information related to disease. However, recent efforts in building more sophisticated machine learning strategies produced unimodal sMRI methods rivaling the state-of-the-art multimodal approaches (Weiner et al. 2017). Although multi-modal approaches tend to be more sensitive in general and offer richer insight, the practical advantages of sMRI being non-invasive, cost-effective and widely-accessible in the clinic, make sMRI-based CAD methods for early prognosis highly desirable.

Cortical thickness is a sensitive imaging biomarker that can be easily derived from sMRI to diagnose AD. However, its sensitivity to identify the prodromal subjects (such as mild cognitive impairment (MCI)) at risk of progressing to AD is limited (Cuingnet et al. 2011). Network-level analysis of cortical thickness and gray matter features demonstrated its potential to provide novel insights or improve predictive power (Raamana et al. 2015), and is gaining in popularity (Evans 2013; Wen, He, and Sachdev 2011; Reid and Evans 2013; Jason P. Lerch et al. 2006). Thickness network features offer complementary information compared to the underlying fiber density (Gong et al. 2012), are shown to be disrupted in AD (Kim et al. 2016) and have been shown to have potential for early prognosis of AD (Raamana et al. 2015; Wee et al. 2012; Dai et al. 2012; Kim et al. 2016), as well as for subtype discrimination (Raamana, Wen, et al. 2014), outperforming the non-network raw-thickness features.

Network analysis studies in cortical thickness range from

1. group-wise studies building networks based on group-wise covariance/correlation in cortical thickness (Evans 2013; He and Chen 2007; Jason P. Lerch et al. 2006), which may be used to characterize the properties of these networks (such as small-worldness) as well as provide useful insight into network-level changes between two diagnostic groups e.g. healthy controls (CN) and Alzheimer’s disease (AD),
2. studies building individual subject-wise graphs based on within-subject ROI-wise (pairwise) similarity metrics (Raamana et al. 2015; Tijms et al. 2012; Wee et al. 2012; Dai et al. 2012; Kim et al. 2016) to enable predictive modeling. These studies resulted in disease-related insights into network-level imaging biomarkers and improved accuracy for the early prognosis of AD. However, these studies are based on distinctly different parcellation schemes of the cortex, vastly different ways of linking two different regions in the brain, and datasets differing in size and demographics.

Insights derived from various brain network studies showed considerable variability in reported group differences (Tijms et al. 2013), and widely accepted standards for network construction are yet to be established (Stam 2014). There have been recent efforts into understanding the importance and impact of graph creation methods, sample sizes and density (van Wijk, Stam, and Daffertshofer 2010; Phillips et al. 2015). However, these studies have been restricted to the choice of group-wise correlation methods to define the edges, or limited to understanding the group-wise differences in selected graph measures. But such important methodological analyses have not been performed in the context of building individual subject-wise predictive modelling. Hence, there is no clear understanding of the impact of different choices in subject-wise network construction and their relative predictive performance.

Given their potential for the development of accurate early prognosis methods (Raamana et al. 2015; Raamana, Wen, et al. 2014) demonstrated by outperforming non-network raw-thickness features, and the wide-accessibility of sMRI, thickness-based networks deserve a systematic study in terms of

1. how does the choice of edge weight or linking criterion (correlation (He and Chen 2007), similarity (Raamana et al. 2015) affect the performance of the predictive models? See Table 3 for more details.
2. how does the scale of parcellation (size and number of cortical ROIs) affect the predictive performance?

These questions, analyzed in our systematic study, can reveal important tradeoffs of this emerging theme of research. In this study, we present a methodological comparison of six different ways of constructing thickness-based, subject-wise networks and present classification results under varying levels of separability. We start with the classic CAD problems i.e. discriminating AD from CN, and mild cognitive impairment (MCI) subjects (converting to AD in ~18 months) from CN in the ADNI dataset. In order to test whether the results from this methodological study generalize to different datasets, diseases and separabilities, we also study the Australian Imaging, Biomarker & Lifestyle Flagship Study of Ageing (AIBL) and the <u>Autism Brain Imaging Data Exchange (ABIDE)</u> datasets. Based on these three large publicly available datasets, we show that the predictive power of single-subject morphometric networks, based on cortical thickness features, is insensitive to spatial scale or edge weight. This is an important finding given we were not only able to replicate these results on an independent dataset, but also replicate them in the presence of a different disease and in a different age group.

## Methods

In this section, we describe the datasets we study in detail, along with a detailed description of the preprocessing and the associated methods.

### ADNI dataset

Data used in the preparation of this article were obtained from the Alzheimer’s Disease Neuroimaging Initiative (ADNI) database (adni.loni.usc.edu). The primary goal of ADNI has been to test whether serial magnetic resonance imaging (MRI), positron emission tomography (PET), other biological markers, and clinical and neuropsychological assessment can be combined to measure the progression of mild cognitive impairment (MCI) and early Alzheimer’s disease (AD). For up-to-date information, see www.adni-info.org.

We downloaded baseline T1 MRI scans (n=671) from the ADNI dataset (Jack et al. 2008), which has quality-controlled Freesurfer parcellation (version 4.3) of the cortical surfaces provided in the ADNI portal (B. Fischl and Dale 2000; Bruce Fischl et al. 2002). The parcellation and cortical thickness values downloaded were carefully visually inspected by the first author PRR for errors in geometry and range. This QC process was rigorous to include a large number of cross-section slices with contours of pial and white surfaces overlaid on the sMRI image in all 3 views. We have also employed external surface views that facilitate easy inspection and identification of any topological defects as well as anatomical accuracy of the Freesurfer labels as a whole. When noticeable errors were found, we eliminated those (n=24) subjects, and no manual editing and corrections were performed. The thickness features from the remaining subjects for the control (CN) and AD groups (effective n=647) comprised the first set of subjects for our analysis in this study. The second set of subjects with a slightly lower level of separability (MCI subjects converting to AD in 18 months, denoted by MCIc) were chosen to match the benchmarking study (Cuingnet et al. 2011) as closely as possible (to enable comparison to the many methods included) based on the availability of their FS parcellation from ADNI and our quality control results. The demographics for the two sets are listed in Table 1.

**TABLE 1:** ADNI I Demographics

| Diagnostic Group | N | Females | Age | MMSE* |
| --- | --- | --- | --- | --- |
| Dataset 1: ADNI |  |  |  |  |
| Healthy controls (CN1) | 224 | 109 | 75.79 (4.99) | 29.11 (1.01) |
| Alzheimer’s disease (AD) | 188 | 89 | 75.22 (7.49) | 23.29 (2.04) |
| Dataset 2: ADNI |  |  |  |  |
| Healthy controls (CN2, for MCI)^ | 159 | 85 | 76.07 (5.33) | 29.17 (0.98) |
| MCI converters to AD in 18 months (MCIc) | 76 | 33 | 74.67 (7.35) | 26.47 (1.86) |

All statistics here are displayed in mean (SD) format.
\*ADNI: Only MMSE is significantly different between CN1 & AD with $p < 0.05$ .
^ADNI: Controls and MCI converters are chosen to match the benchmark samples presented in (Cuingnet et al. 2011) as closely as possible allowing for exclusions due to quality control.

### AIBL Dataset

In order to study whether the results from the ADNI dataset in this study generalize to another independent dataset, we have downloaded the Australian Imaging, Biomarker & Lifestyle Flagship Study of Ageing (AIBL) dataset (Ellis et al. 2009), which contained similar (but not identical) patient groups and diagnostic categories. The downloaded subjects were processed with Freesurfer v6.0. The number of Alzheimer’s subjects we could download from AIBL (denoted by AD2) at baseline were n=64, and we randomly selected 100 healthy controls (CN4) for this study, whose subject IDs are shared in the Appendix. The resulting cortical parcellations were visually quality controlled by PRR with VisualQC (v0.4.1) (Raamana 2018; Raamana and Strother 2018b). This QC process was rigorous to include a large number of cross-sectional slices with contours of pial and white surfaces overlaid on the sMRI image in all 3 views, with the ability to zoom in to the voxel-level to ensure anatomical accuracy of the pial and white surfaces. In addition, the VisualQC interface presents 6 views of the external surface view of pial surface which facilitates easy inspection and identification of any topological defects as well as anatomical accuracy of the Freesurfer labels as a whole. This QC process was employed to remove subjects with inaccurate parcellations or any other errors that render them unusable for analyses in this study. We would like to note that VisualQC is the most comprehensive QC tool for Freesurfer parcellations, and hence this process may be sensitive to catching the parcellation errors compared to that on the non-interactive tool employed on the ADNI dataset. This resulted in a usable subset of 51 AD2 and 80 CN3 subjects. Demographics of the subjects analyzed are presented in Table 2.

**TABLE 2:** AIBL Demographics

| Diagnostic Group | N | Females | Age | MMSE* |
| --- | --- | --- | --- | --- |
| Healthy controls (CN4) | 80 | 44 | 73 (6.99) | 29 (1.26) |
| Alzheimer’s disease (AD2) | 51 | 32 | 73 (7.37) | 22 (5.4) |

All statistics here are displayed in median (SD) format.

Motivated by intention to improve reproducibility and maximize the value of this study by employing open source tools when possible, we have employed graynet (Raamana and Strother 2018a, 2017) to compute the network-level features, and neuropredict (Raamana 2017) to evaluate their predictive utility. While this change of software libraries would add another interesting level of robustness check for the results presented here, we must note that this change in technology stack may lead to some differences in numerical estimates e.g. in AUC estimates when comparing across different datasets e.g. ADNI vs. AIBL. However, given the technology employed is the same for a given dataset, the performance estimates within the dataset will be perfectly commensurable for posthoc statistical analyses.

### ABIDE dataset

In order to study whether the conclusions drawn from the ADNI dataset generalize to a very different disease cohort, we obtained the Freesurfer parcellations (version 5.1) from the <u>Autism Brain Imaging Data Exchange (ABIDE)</u> preprocessed dataset made available freely on the <u>ABIDE website</u> (Craddock, Cameron and Benhajali, Yassine and Chu, Carlton and Chouinard, Francois and Evans, Alan and Jakab, Andr?s and Khundrakpam, Budhachandra Singh and Lewis, John David and Li, Qingyang and Milham, Michael and Yan, Chaogan and Bellec, Pierre 2013). A random subset of cortical parcellations (n=227) have been visually inspected by PRR for errors in geometry estimation and value ranges (using the same in-house tools and processed used on the ADNI dataset) to eliminate any subjects showing even a mild chance of failure. From the passing subjects (n=226), we randomly selected 200 subjects (100 samples per diagnostic group) whose demographics are presented in Table 3 and the subject IDs are listed in the Appendix. Owing to the random selection, they come from multiple sites, which is akin to the ADNI dataset used in this study. Previous research (Abraham et al. 2017) showed that the site heterogeneity has little or no impact on the predictive accuracy of network-level features derived from task-free fMRI data. The distribution of the sites represented in this study are shown in Appendix D.

**Table 3:** ABIDE I demographics

| Dataset 3: ABIDE |  |  |  |  |  |  |
| --- | --- | --- | --- | --- | --- | --- |
| Diagnostic Group | N | Females | Age | FIQ* | PIQ | VIQ* |
| Healthy controls (CN3) | 100 | 17 | 17.27 (7.68) | 109.10<br>(12.35) | 105.64<br>(12.74) | 111.89 (13.52) |
| Autism (AUT) | 100 | 9 | 15.82 (5.93) | 103.49<br>(14.68) | 104.57<br>(14.68) | 101.36 (15.86) |
| ABIDE: FIQ and VIQ are significantly different between CN3 & AUT with $p < 0.05$.<br>FIQ: Full IQ standard score<br>VIQ: Verbal IQ standard score<br>PIQ: Performance IQ standard score | | | | | | |

### Feature extraction

In the following sections, we describe the steps involved in the extraction of weighted networks based on T1 MRI scans of the different subjects in the two independent datasets.

#### Alignment and dimensionality reduction

Cortical thickness features studied here were obtained from the Freesurfer parcellations (gray and white matter surfaces). They were then resampled to the *fsaverage* atlas and smoothed at *fwhm*=10mm to reduce the impact of noise. This is achieved by Freesurfer ‘-qcache’ processing option, which registers each of the subjects to the *fsaverage* atlas (provided with Freesurfer) to establish vertex-wise correspondence across all the subjects.

#### Cortical subdivision

In order to avoid the curse of dimensionality and to reduce the computational burden, the atlas has been subdivided using a surface-based, patch-wise parcellation technique originally presented in (Raamana et al. 2015). This technique is based on Freesurfer parcellation which consists of 34 ROIs per hemisphere, which vary in size (number of vertices) greatly. In order to obtain uniform sized patches, we subdivide each of these ROIs into smaller patches, while respecting the anatomical boundaries of each ROI. Here, we use an adaptive version wherein the patch-size is controlled by number of vertices (denoted by *m*=vertices/patch), instead of choosing a globally fixed number of patches (say 10) per Freesurfer APARC label regardless of its size (which can vary widely resulting in vastly different patch sizes within the same subject). As we change *m*, the subdivision of the cortical labels is performed purely on the existing mesh, and neither the geometrical parcellation itself nor the vertex density are modified. Here, *m* can be taken as the size of the graph node (imagine the node as a small patch within different Freesurfer labels). Alternatively, *m* can be seen as the spatial scale of the graph analysis, whose impact is being assessed for different values of *m*. When m is small (say 100), this results in large number (273) of total patches (sum of number of patches for each aparc label) across the whole cortex, whereas it results in only 68 patches when it is very high (m=10000), as such a large patch covered the full extent of all the 68 Freesurfer APARC labels currently defined on fsaverage cortical parcellation. We have analyzed the following values of *m*= 1000, 2000, 3000, 5000 and 10000, which resulted in the following total number of non-overlapping patches in the whole cortex: 273, 136, 97, 74 and 68 respectively.

**Table 3:** Variety of edge weights analyzed in this study.

| Type of base representation | Type of edge weight metric | Acronym | Mathematical definition |
| --- | --- | --- | --- |
| Summarized<br>(median/ mean in<br>a patch) | Similarity<br>(Raamana et al. 2015) | MD | $\vert M_i - M_j\vert $ |
| | exp(similarity) | EMD | $e^{-\frac{(\mu_i - \mu_j)^2}{2(\sigma_i + \sigma_j)}}$ |
| Raw distribution | Wilcoxon ranksum<br>statistic | RS | Ranksum test statistic |
| Normalized<br>histogram | Histogram correlation | HCOR | $\rho(h_i, h_j)$ |
| | $\chi^2$ statistic | CHI2 | $2 \sum_{k=1}^N \frac{(h_i(k) - h_j(k))^2}{h_i(k) + h_j(k)}$ |
| | Histogram intersection | HINT | $\frac{\sum_{k=1}^N \min(h_i(k), h_j(k))}{\sum_{k=1}^N h_i(k)}$ |

*This table presents the list of edge weights compared in this study. Notation: each patch is indexed by $i$ or $j$ . For patches $i$ and $j$ , $M$ , $\mu$ and $\sigma$ are the median, mean and standard deviation of the within-patch distribution of vertex-wise thickness values; $h_i$ is the normalized histogram of a given distribution. $N$ is the number of bins in the histogram, which is fixed at $N=100$ bins.; $\rho$ is the Pearson correlation coefficient between two vectors of equal length.*

#### Network Computation

Construction of thickness networks in their early form were based on group-wise correlations (He and Chen 2007). Our previous publications based on cortical thickness (Raamana et al. 2015; Raamana, Wen, et al. 2014) and other interesting studies on gray matter density (Tijms et al. 2012; Wee et al. 2012) extend the earlier approaches to individualized subject-wise network extraction methods. Many of these previous studies relied on summarizing the thickness distribution in a given ROI (e.g. using mean within the entire Freesurfer label as in (Tijms et al. 2012)) or within a patch (Freesurfer label subdivided further as in (Raamana et al. 2015)), before constructing the networks. Although such approaches reduce the dimensionality and provide us with smooth features, they do not utilize the rich description and variance of the distribution of features. Moreover, studies thus far computed characteristic features from a binary network (by applying an optimized threshold (Raamana et al. 2015)) or using a vector representation of weighted graphs (vector of distances in the upper triangular part of the edge weight matrix, as they are symmetric (Tijms et al. 2012)). Here, in order to enable a principled comparison across the different edge weights (and to avoid the optimization of an arbitrary threshold required to binarize the edge weight matrix), we study weighted-networks only, whose derivation is described below.

#### HIstogram WEighted NETworks (HiWeNet)

In this section, we describe the method employed in constructing the HIstogram WEighted NETworks (HiWeNet) based on cortical thickness. First, to improve the robustness of the features, 5% outliers from both tails of the distribution of cortical thickness values are discarded from each patch at a given scale *m* (see Appendix for more information). The residual distribution is converted into a histogram by binning into uniformly spaced *n* = 100 bins. Then the histogram counts are normalized using

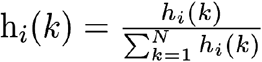

for *k* = 1 : *N*, where *h_i_* is the histogram of patch *i*. This method (illustrated further in Figure 1) enables the computation of the pairwise edge-weight (distance between the histograms, denoted by EW) for the two patches *i* and *j*. A variety of histogram distances as listed in Table 3 are studied in this paper to analyze their impact on predictive power.

**Fig. 1:**
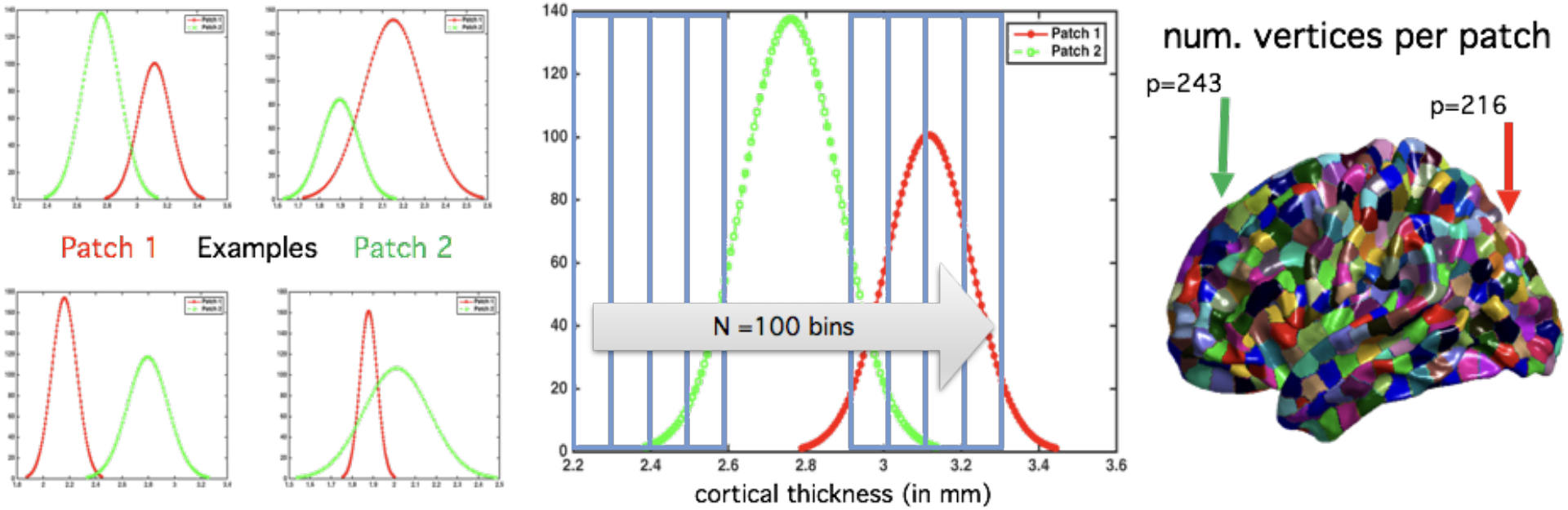
Construction of histogram-distance weighted networks (HiWeNet) based on cortical thickness features using edge-weight calculations (applicable to HCOR, CHI2 and HINT metrics in Table 3). The four smaller subpanels on the left show typical distributions of cortical thickness values for four random pairs of patches (in green and red) in a given subject (shown on cortical visualization on right). They demonstrate the means and shape of these distributions can vary substantially as you traverse across different pairs of cortical patches. The large panel in the middle illustrates the type of binning used to construct the histogram from each patch.

To analyze the relative benefit of HiWeNet, we compare the histogram-based methods to three commonly used inter-nodal weights based on descriptive summary statistics (denoted as MD, EMD and RS in Table 3). Once the edge weight matrix is computed (which is symmetric), we extract the upper-triangular part of the matrix and vectorize it (of length *n*(n-1)/2*, where *n* is the number of patches on the cortex for a given number of vertices/patch *m*). The vectorized array of edge weights (VEW) forms the input to the classifier. Each element of VEW corresponds to a unique edge in the matrix of pairwise edges. In addition, in Appendix C, we present and analyze the performance of an alternative network-representation method.

#### Note on test-retest reliability

The reliability of this network approach developed in *HiWeNet* (pairwise distances between ROIs) boils down to the reliability of the method to measure cortical thickness at the vertex-level, as the remaining parts of the algorithm are deterministic. Several studies have previously shown that cortical thickness estimation (and Freesurfer as a tool) have high test-retest reliability (Han et al. 2006; Iscan et al. 2015) and that the brain-behaviour relationships e.g. between cortical thickness and cognitive performance are stable across different sessions, scanner platforms and field strengths (Dickerson et al. 2008). In addition, given our choice of employing distance between thickness distributions over relatively large patches (1000 vertices or more), small changes in thickness (e.g. 0.2mm) would be absorbed into the distance calculations, and hence are unlikely to change the results presented herein.

#### Comparison of predictive utility

In this section, we describe the procedure and techniques used to evaluate and compare the predictive power of multiple variations of the network-level features. Thanks to the relatively large sample sizes, particularly for ADNI and ABIDE, we could employ a repeated nested split-half cross-validation (CV) scheme, with 50% reserved for training, in order to maximize the sizes of training and test sets. Moreover, in each iteration of CV, all the methods are trained and assessed on the exact same training and test sets, in order to “pair” the performance estimates. This technique is shown to produce reliable and stable estimates of differences in predictive performance across different methods (Dietterich 1998; Burman 1989; Demšar 2006), instead of pooling multiple sets of performance distributions estimated separately on different training and test sets for each method independently. This setup allows us to compare large numbers of methods and their variants simultaneously within each dataset.

##### Cross-validation scheme

The comparison scheme employed is comprised of the following steps – for a schematic, see Fig. 4 in (Raamana et al. 2015):

1. repeated split-half cross-validation scheme, with class-sizes stratified in the training set (RHsT) (Raamana et al. 2015), to minimize class-imbalance. This scheme is repeated N=200 times, to obtain the N paired estimates of classification performance.
2. In each CV run,

a. feature selection (from vectorized array of edge weights, VEW) on one split (training set of size N_train_) is performed based on t-statistic based ranking (based on group-wise differences in the training set only), selecting only the top N_train_/10 elements. The frequency of selection of a particular element (which is an edge in the cortical space) over different CV trials by the t-statistic ranking is an indication of its discriminative utility, and will be visualized to obtain better insight into the process.
b. Support vector machine (SVM) is chosen as the classifier to discriminate the two groups in each experiment. SVM is optimized in an additional inner split-half CV applied to the training set via a grid search. We have employed the following ranges of values in the grid search for the margin control parameter C = 10p; p = - 3 : 5 and the kernel bandwidth = 2q; q = −5 : 4.
c. The optimized SVM is tested on the second split (test) to evaluate its performance.
3. The process in Step 2 is repeated N=200 times (Varoquaux et al. 2016; Raamana et al. 2015) to obtain 200 independent estimates for each method being compared.
4. In this study, we measure the performance by area under the predictive receiver operating characteristic (ROC) curve (denoted by AUC), whose distributions for different methods are shown in Figure 3.

The results in this study were produced using Raamana’s programming library implemented in Matlab based on the built-in statistics and machine learning toolbox.

#### Open source software

Most of the computational code applied on the ADNI and ABIDE datasets had been implemented in Matlab. In order to enable other researchers to utilize the methods presented here easily without having to pay for expensive Matlab licenses, we have re-implemented them in python following the best practices of open science. Moreover, we have processed the AIBL dataset using the open source alternatives, and showed that our results replicate on an independent dataset, despite differences in the following 3 layers of software.

##### Quality Control via VisualQC

To achieve higher rigor as well as ease of use, we have developed an interactive version of Freesurfer QC tool, which is available as part of the VisualQC package at github.com/raamana/VisualQC. This tool, applied on AIBL dataset, is more sensitive in detecting parcellation errors compared to the in-house Matlab tools and other existing protocols applied on the ADNI and ABIDE datasets (study to be published).

##### Feature Extraction via graynet

The core HiWeNet algorithm has been implemented in Python and is publicly available at this URL: https://github.com/raamana/hiwenet (Raamana and Strother 2017). We have also published the original Matlab code for the computation of adjacency matrices used for this study, within the *hiwenet* package.

Further, in order to make this research even more accessible, we have implemented the entire workflow of morphometric network extraction as a seamless pipeline called *graynet*, implemented entirely in Python (Raamana and Strother 2018a). Using this tool would enable those without much software engineering experience to simply run Freesurfer and then run *graynet* to get started with morphometric network analyses. This frees them from the hassle of assembling complicated data, implementing graph theoretical operations and managing the pipeline following the best practices, which can be a barrier to many laboratories with limited computational and software expertise. In addition, we employed this tool to process the AIBL dataset, and show that patterns in performance comparison across different weights are retained compared to those of the original Matlab toolbox.

##### Evaluating predictive utility via neuropredict

In order to enable a much wider audience (those without access to a Matlab license or its expensive statistical learning toolboxes (each to, or those who do not have the necessary programming skills or machine learning expertise) utilize a comprehensive performance evaluation tool, we have also built an open source tool called *neuropredict* (Raamana 2017) at github.com/raamana/neuropredict. Once the researchers run Freesurfer successfully, they can run *graynet* (Raamana and Strother 2018a), which produces the necessary single-subject morphometric networks. The outputs from graynet in turn serve as direct input to neuropredict, which runs the cross-validation scheme described in the above section to produce a comprehensive report on their predictive power. In addition, we employed this tool to evaluate the performance of network features from the AIBL dataset (CN4 vs. AD2), matching the techniques and specific optimizations to the extent possible. This showed that our findings replicated compared to that of the original Matlab toolbox, which validates neuropredict as a useful open source alternative.

## Results and Discussion

### Within-group networks

To obtain better insight into the topology of the networks defined above, it is helpful to visualize seed-based networks and analyze their connections. A common approach to this end involves picking the posterior cingulate gyrus (core hub of the default mode network, DMN) as the seed and analyzing its connections in healthy controls, and esp. how they change for different edge metrics. The seed-based network visualizations are produced for each edge weight method separately for m=2000, identical to the network construction method described in the Methods section: compute histogram-distance between the thickness distribution of the seed and all the other ROIs, averaging this edge weight across all the healthy subjects, and retaining only the strongest edges (top 5%).

To make the comparison across the three datasets easy, they are grouped for each metric e.g. for median difference (MD), the comparison is shown below for healthy controls. From this figure, we can clearly see a pattern resembling the default mode network, in healthy controls from all the three samples. This is consistent with the results reported in previous structural covariance studies (Spreng and Turner 2013; Evans 2013; Spreng et al. 2013; Power et al. 2011).

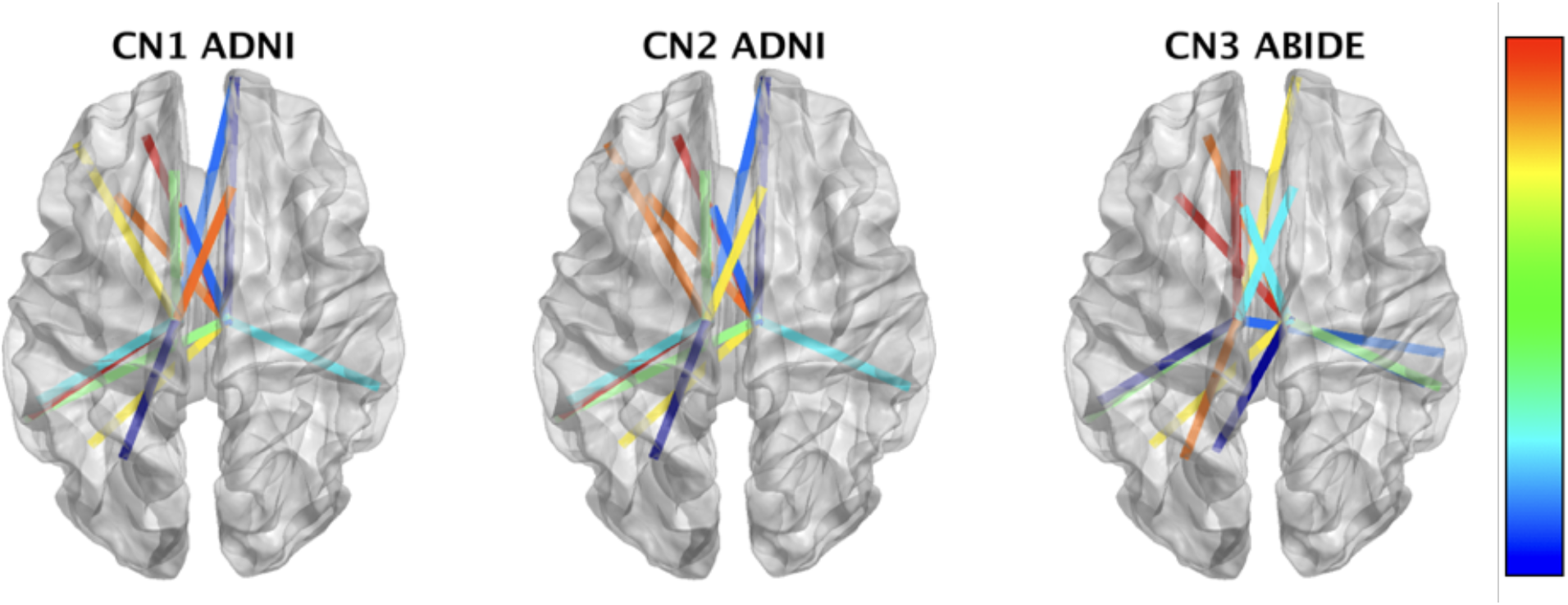
Caption: seed-based connectivity networks for the MD metric (m=2000), showing average weights across each healthy control sample from the three datasets (as labelled). The colors on the edges represent the edge weight using a jet colormap (with blues indicating the weaker and reds indicating stronger weights). From this figure, we can clearly see a pattern resembling the default mode network, in healthy controls from all the three samples.

To get a sense of how these networks change with different EW metric, we show two other networks corresponding to HCOR and CHI2 metrics below (each figure is labelled with the metric and summary statistic being displayed e.g. HCOR mean). This HCOR network loses resemblance to the DMN (e.g. loss of edges to superior frontal, banks of the superior temporal sulcus, frontal pole, fusiform), and the edge weight distribution varies widely across the three samples. However, the CHI2 network resembles the DMN pattern seen in MD network well, suggesting the similarity of the two networks.

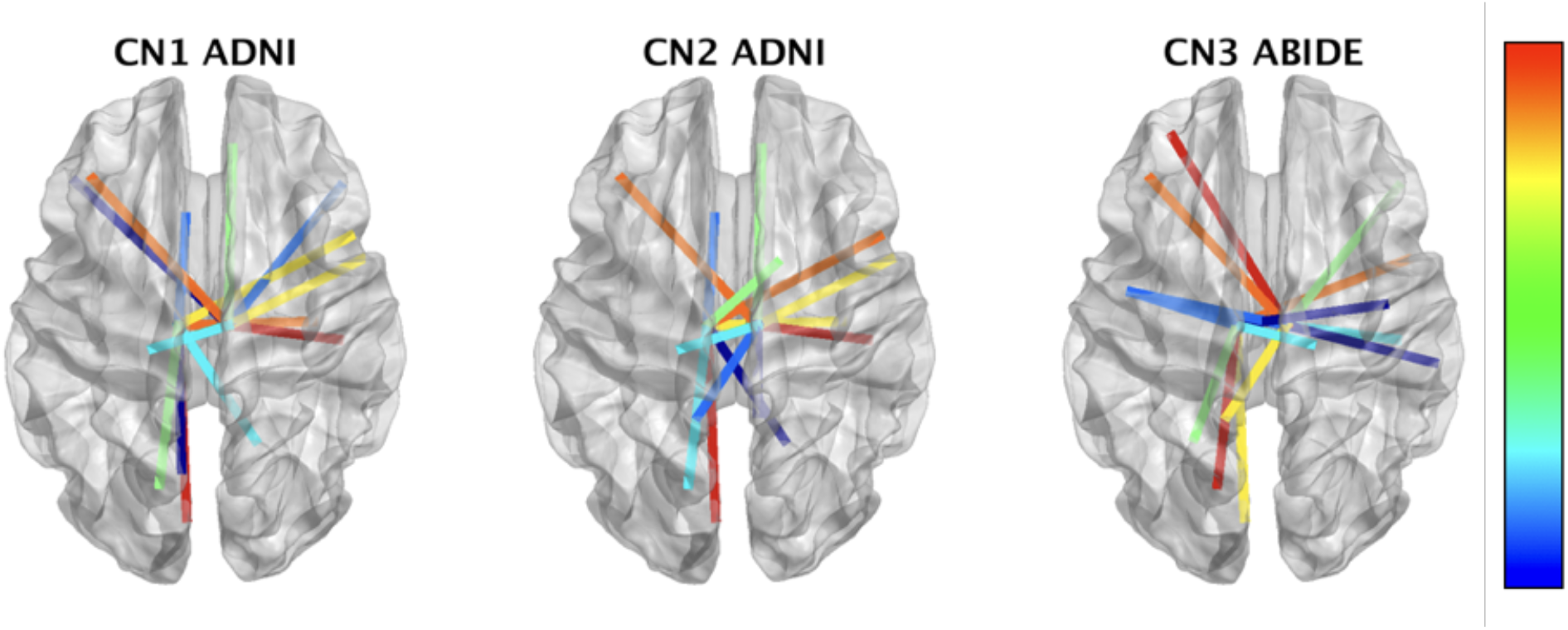
Caption: network showing edge weights(mean across samples) derived via HCOR metric. Layout of the figure is the same as above for the MD network.

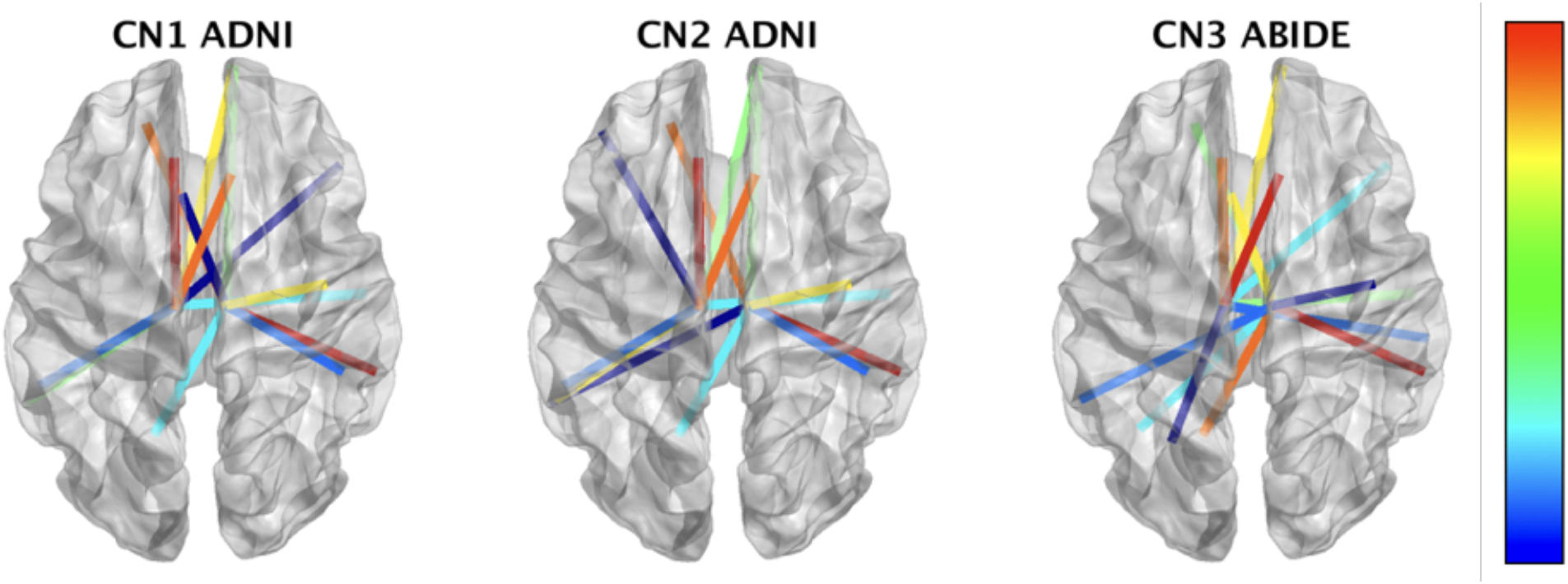
Caption: network showing edge weights (mean across samples) derived via CHI2 metric. Layout of the figure is the same as above for MD.

### Group-wise differences

To illustrate the differences between the proposed methods of computing edge weights, we compute the distributions of vertex-wise mean thickness values for CN1 and AD separately. We then visualize them in the form of a matrix of pairwise edge weights at *m*=2000, as shown in Figures 2 (a) and 2(b). Each row (say node *i*) in a given edge-weight matrix (from one group say CN1 in Fig. 2 (a)) here refers to the pairwise edge weights w.r.t remaining nodes *j, j = 1:*N. As the differences are subtle and spatially distributed, for easy comparison between the two classes, we visualize the arithmetic differences between the two classes in Fig. 2 (c).

**Fig. 2.**
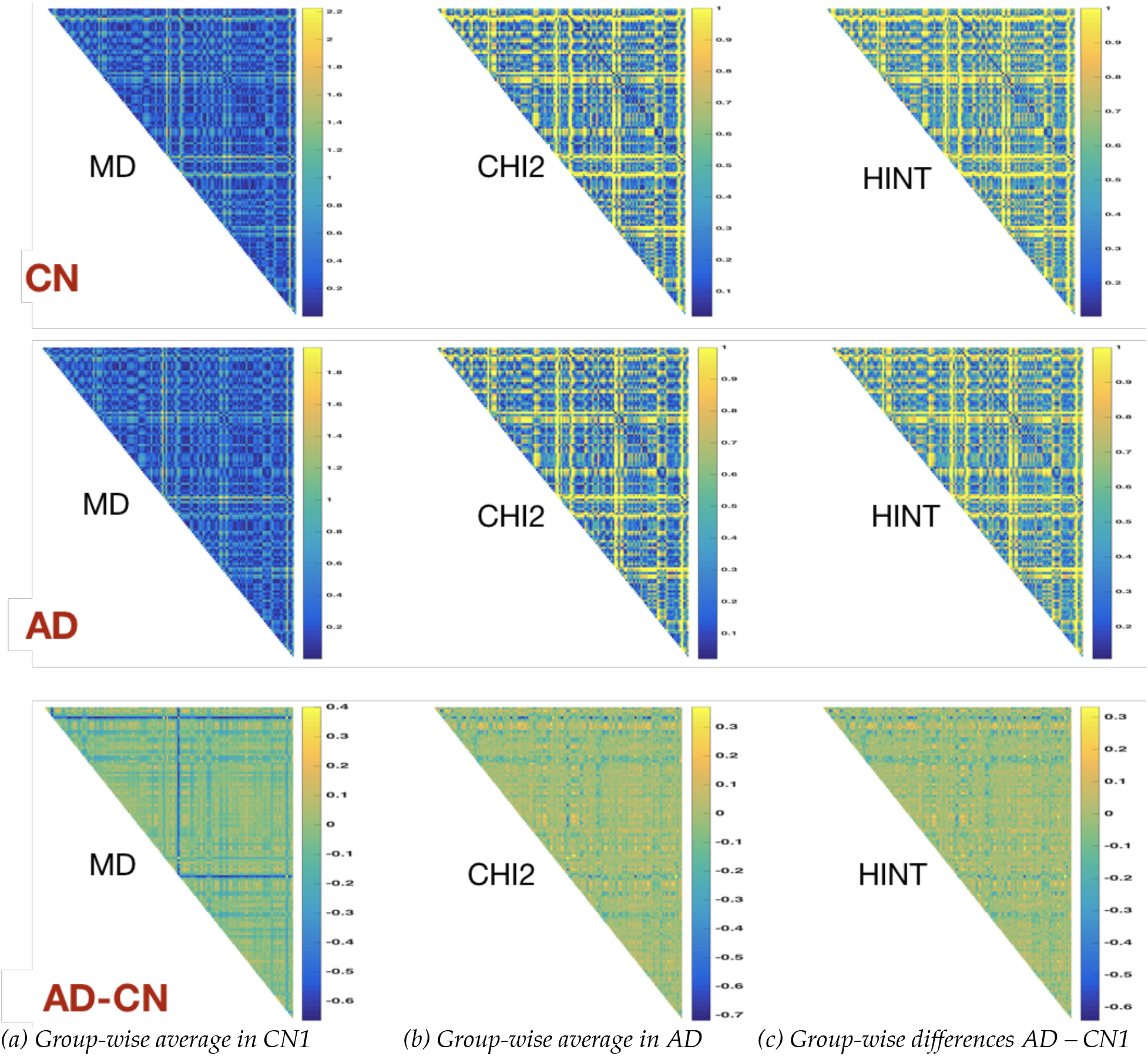
Edge weights derived from group-wise average thicknesses for three definitions of edge weight. (a) healthy controls (CN1) (b) Edge weights group-wise average in Alzheimer’s disease (AD), both at m=2000. (c) Arithmetic differences i.e. AD – CN1. The three panels in each subfigure show the edge weights from MD, CHI2 and HINT methods as defined in Table 3. In each of the panels, we present the upper triangular part of the edge-weight matrix (pairwise) computed using the corresponding equations in Table 3. We notice there are clear differences among the patterns in the three panels. The panels (a) and (b) appear similar at first glance, but they are sufficiently different to be observed in panel (c).

The visualizations in Fig. 2(c) offer useful insight into the group-wise differences between CN1 and AD, and across different edge weight distances. However, visual differences do not imply differences in predictive power of features extracted these networks of weights. Hence, it is important to assess their predictive utility in discriminating AD from CN1.

### Predictive utility

The RHsT cross-validation scheme is employed for each of the three classification experiments from two independent datasets i.e. CN1 vs. AD, CN2 vs. MCIc and CN3 vs. AUT. The performance distributions for the different combinations are shown in Fig. 3.

**Fig. 3.**
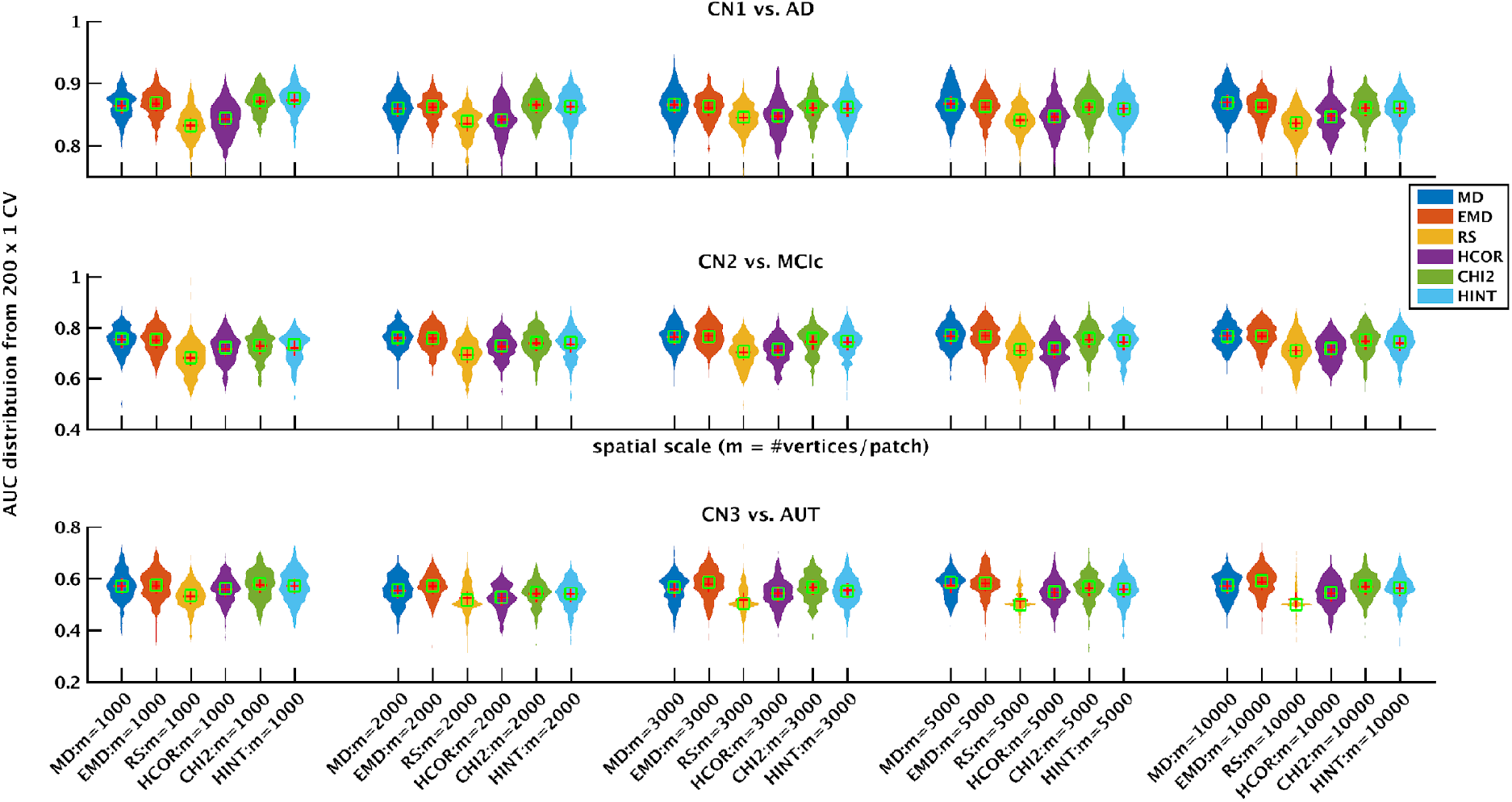
Classification performance for the different network methods (different edge weight metrics at different spatial resolutions of m) in discriminating AD (top panel), MCIc (middle panel) and AUT (bottom) panel from their respective control groups under a rigorous CV scheme. The data for three experits come from ADNI, ADNI and ABIDE datasets respectively (see Tables 1 and 2). The performance presented here is a distribution of AUC values from 200 randomized train/test splits of RHsT (whose median is shown with a red cross-hair symbol).

Focusing on the top panel (CN1 vs. AD), there are numerical differences in performance among different methods at fixed scale (*m*). However, the pattern remains similar across different spatial scales. The MD, EMD, CHI2 and HINT methods are consistently outperforming, numerically speaking, the RS and HCOR methods across different values of *m*. Broadly speaking, the patterns of change in AUC in Fig. 3 within each panel as we move from left to right (going over different combinations) are quite similar to the rest, although at a different median baseline (at AUC=0.87 for CN1 vs. AD, at AUC=0.75 for CN2 vs. MCIc and at AUC=0.6 for CN3 vs. AUT).

### Statistical significance testing

In order to assess the statistical significance of differences among this large set of methods, we performed a nonparametric Friedman test (Dietterich 1998) comparing the performance of the 30 different classifiers (6 methods at 5 spatial scales) simultaneously, for each of the three experiments separately. The results from post-hoc Nemenyi test (Demšar 2006) are visualized in a convenient critical difference (CD) diagram (Kourentzes 2016) as shown in Figure 4.

**Fig. 4.**
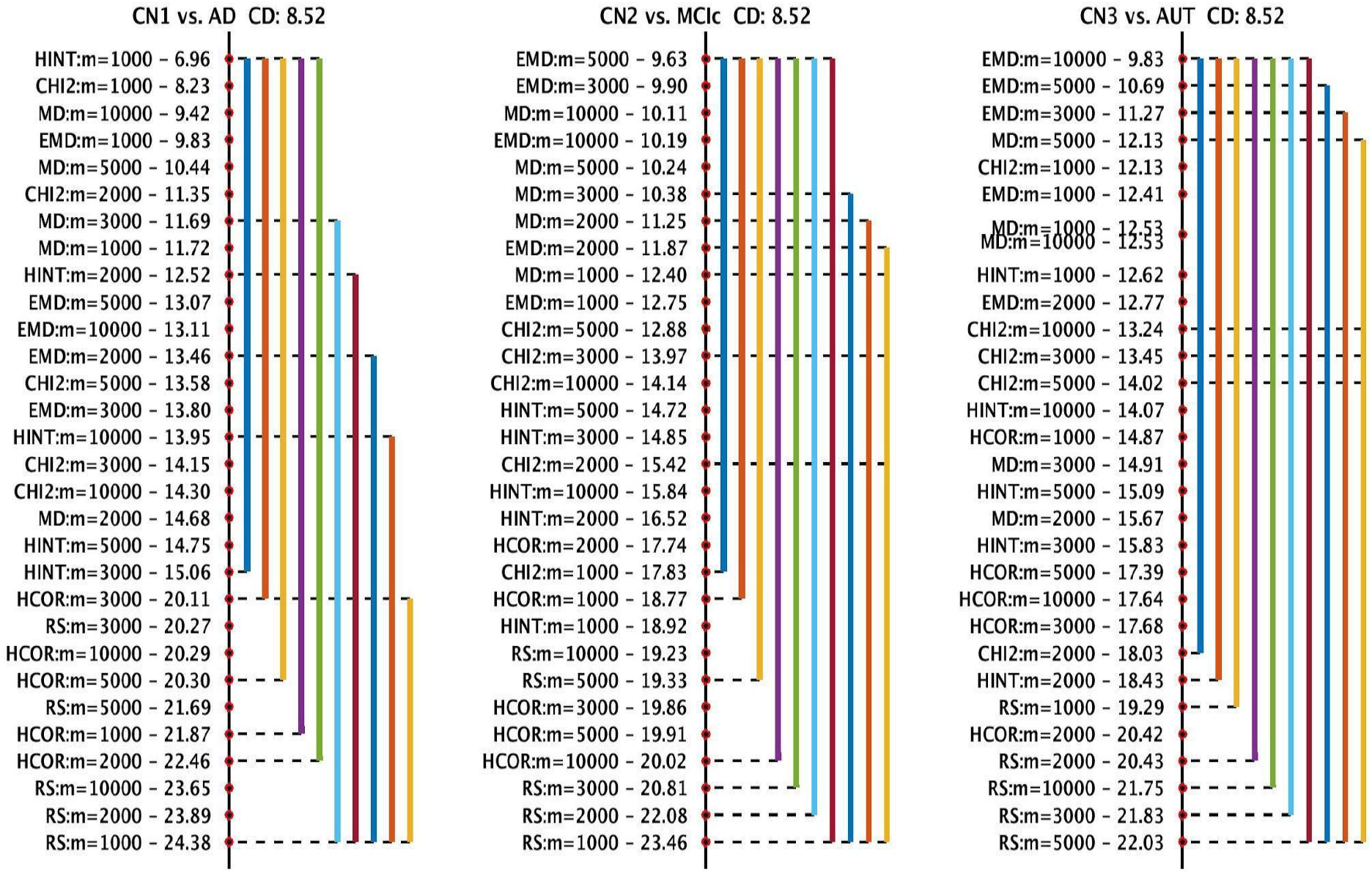
Critical difference diagram comparing the ranks of different classification methods in a non-parametric Friedman test based on classification performance results from a rigorous CV evaluation method using 200 iterations of holdout. Here, smaller numerical values for rank implies higher performance. The vertical axis presents the ranks (better ranks and methods at the top, and worse ranks and methods to the bottom). The performance of any two methods are statistically significantly different from each other, if their ranks differ by at least the critical difference (CD), which is noted on top of each of the three panels. If a group of methods (annotations on the left within each panel) are connected by a line, they are not statistically significantly different from each other. Different colored lines here present groups of methods that are not significantly different from each other in ranks, each one using a different method as its reference point. For example, in the leftmost panel presenting the results from CN1 vs. AD experiment, the leftmost blue line connects all the methods between the highest ranked HINT:m=1000 (ranked 6.96) to the HINT:m=3000 method (ranked 15.06), including themselves, which implies they are not statistically significantly different from each other. In the same panel, the highest-ranked HINT:m=1000 method is not connected to RS:m=1000 (least-ranked 24.38) via any of the colored lines - hence they are indeed statistically significantly different from each other (difference in ranks higher than CD). The values of m= 1000, 2000, 3000, 5000 and 10000 correspond to the following total number of non-overlapping patches in the whole cortex: 273, 136, 97, 74 and 68 respectively

The left panel in Fig. 4 shows that only the top 6 methods (with median ranks from 6.96 to 11.35) are statistically significantly different from the lowest-ranked methods, at *α*=0.05, correcting for multiple comparisons. The remaining 24 methods, when compared together simultaneously, are not significantly different from each other. Similarly, the top-ranked 6 methods are not statistically significantly different from each other. We observe a similar pattern in the center panel (CN2 vs. MCIc), except only the top 5 are statistically significantly different from the lowest ranked methods. In the CN3 vs. AUT case, there are no significant differences at all, possibly due to rather low performance from all the methods to begin with (median AUC across methods is around 0.55).

When the comparison is made at a fixed scale *m*, within each experiment, the performance of the 6 different methods (simultaneous comparison of 6 methods) for most values of *m* are not statistically significantly different from each other, except for m=1000 (CD diagrams are not shown). When the comparison is done for a fixed edge-weight metric at different values of *m*, the performance is not statistically significantly different for any m. Also, the top 2 methods are MD and EMD networks (based on differences in median and mean respectively) at the highest resolution m = 1000 and also at the lowest resolution m = 10,000. This indicates that impact of the nodal size on the predictive performance of a network method may be insignificant. This result is consistent with the findings of (Zalesky et al. 2010; Evans 2013), wherein it was observed that group-wise small-worldness and scale-freeness are unaffected by spatial scale.

### Most discriminative regions

As noted in our CV section earlier, our method records the frequency (across the N CV iterations) of selection (of each weighted connection in VEW) from the t-statistic based ranking method applied on the training set. This helps us gain insight into which pair-wise links have been most frequently discriminative. This pair-wise link frequency can be mapped back to individual cortical patches for intuitive visualization, identifying most discriminative regions (MDRs). One such visualization, thresholding the importance at 50% derived at *m*=2000, is shown in Fig 5. Each color of patch on the cortex represents a particular EW metric (labelled on the colorbar) that led to its selection, and when multiple methods selected the same region (indicating additional importance), we painted it red and labelled it “Multiple”. Note the input to the SVM classifier was a vector of edge weights (from upper-triangular part of the edge weight matrix), and hence the selection of a particular edge leads to highlighting both the regions forming the link. Moreover, the importance of a particular node (cortical patch) could be accumulated from its multiple links, if any.

**Fig 5:**
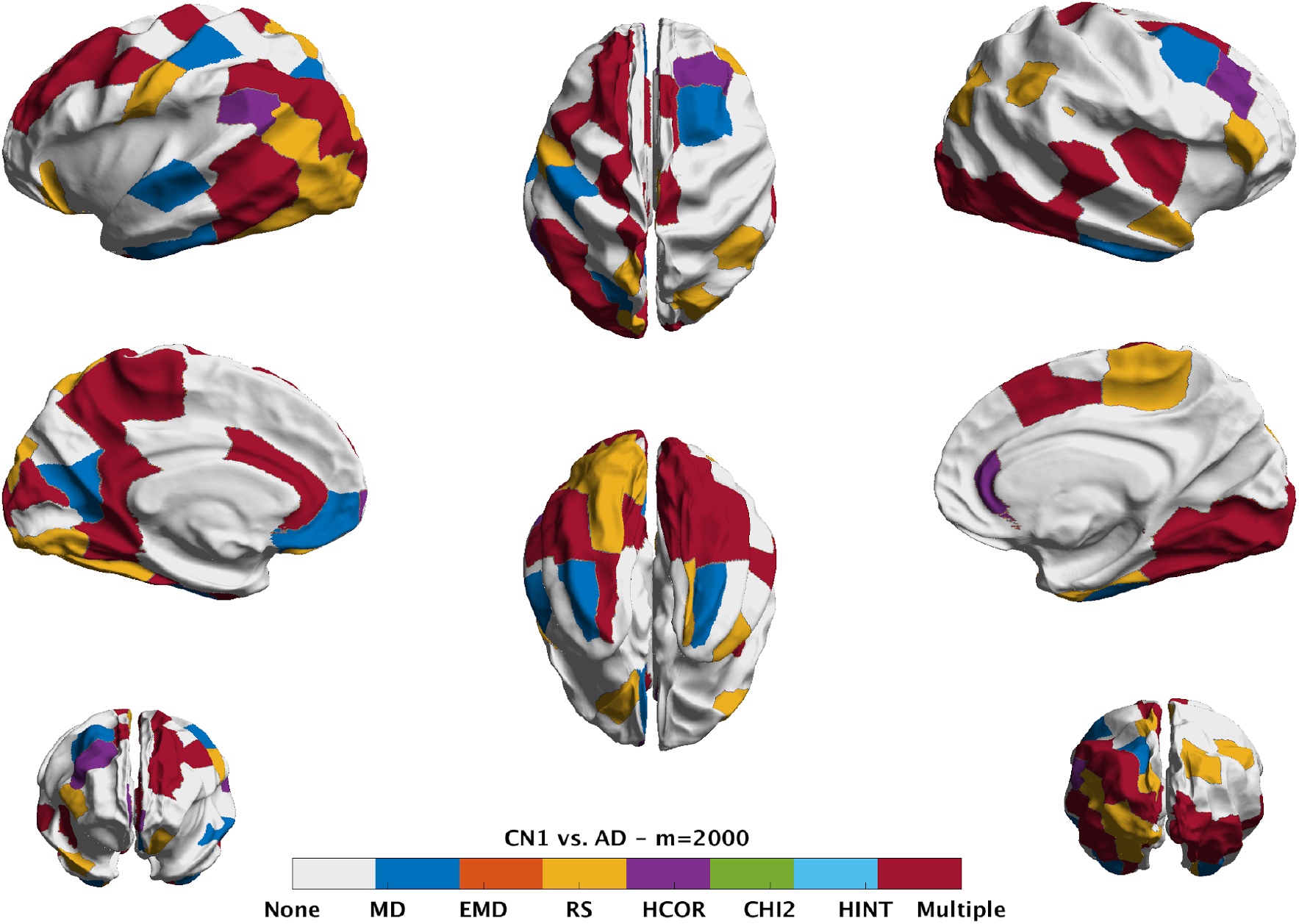
Visualization of the most discriminative regions as derived from the CN1 vs. AD experiment at m=2000. Due to the distributed nature of the degeneration caused by AD, we expect the MDRs to span a wide area of the cortex as observed here. The color of the patch on the cortex represents a particular EW metric (labelled on the colorbar) that led to its selection, and when multiple methods selected the same region (indicating additional importance), we painted it red and labelled it “Multiple”.

Fig. 5 shows the red MDRs (identified by multiple methods as MDR) cover a large cortical area, which is not unexpected, given the changes caused by full AD are known to be widespread over the cortex. In Fig. 6, we observe the MDRs in areas consistently identified with progressive MCI or early stage AD such as middle temporal lobe, cingulate (anterior and inferior), cuneus and precuneus. Of interest here is the clear hemispheric asymmetry to the left, which can also be observed to a lesser extent in the MDRs for AD in Fig. 5. The MDRs identified in discriminating AUT from CN3 are shown in Fig. 7. They appear in the lingual, supra-marginal, post- and precentral areas, which are consistent with previous reports on Autism studying the group differences in developmental patterns of cortical thickness (Smith et al. 2016; Scheel et al. 2011), as well as found to be important in other prediction tasks (Moradi et al. 2017).

**Fig 6:**
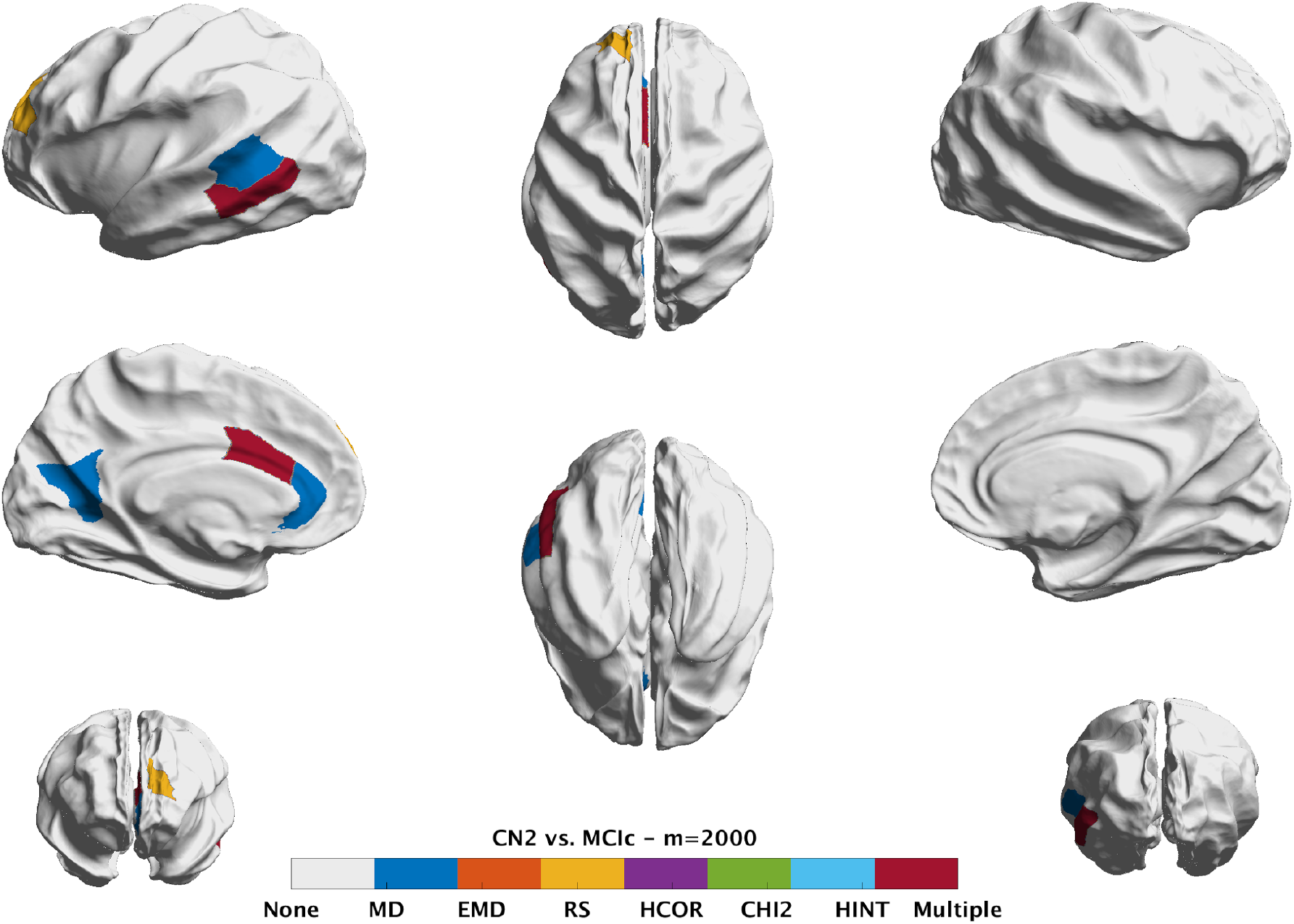
Visualization of the most discriminative regions as derived from the CN2 vs. MCIc experiment at m=2000. MDRs in this experiment identify regions in middle temporal lobe, cingulate (anterior and inferior), cuneus and precuneus, which are known to be associated with progressive MCI and prodromal AD.

**Fig 7:**
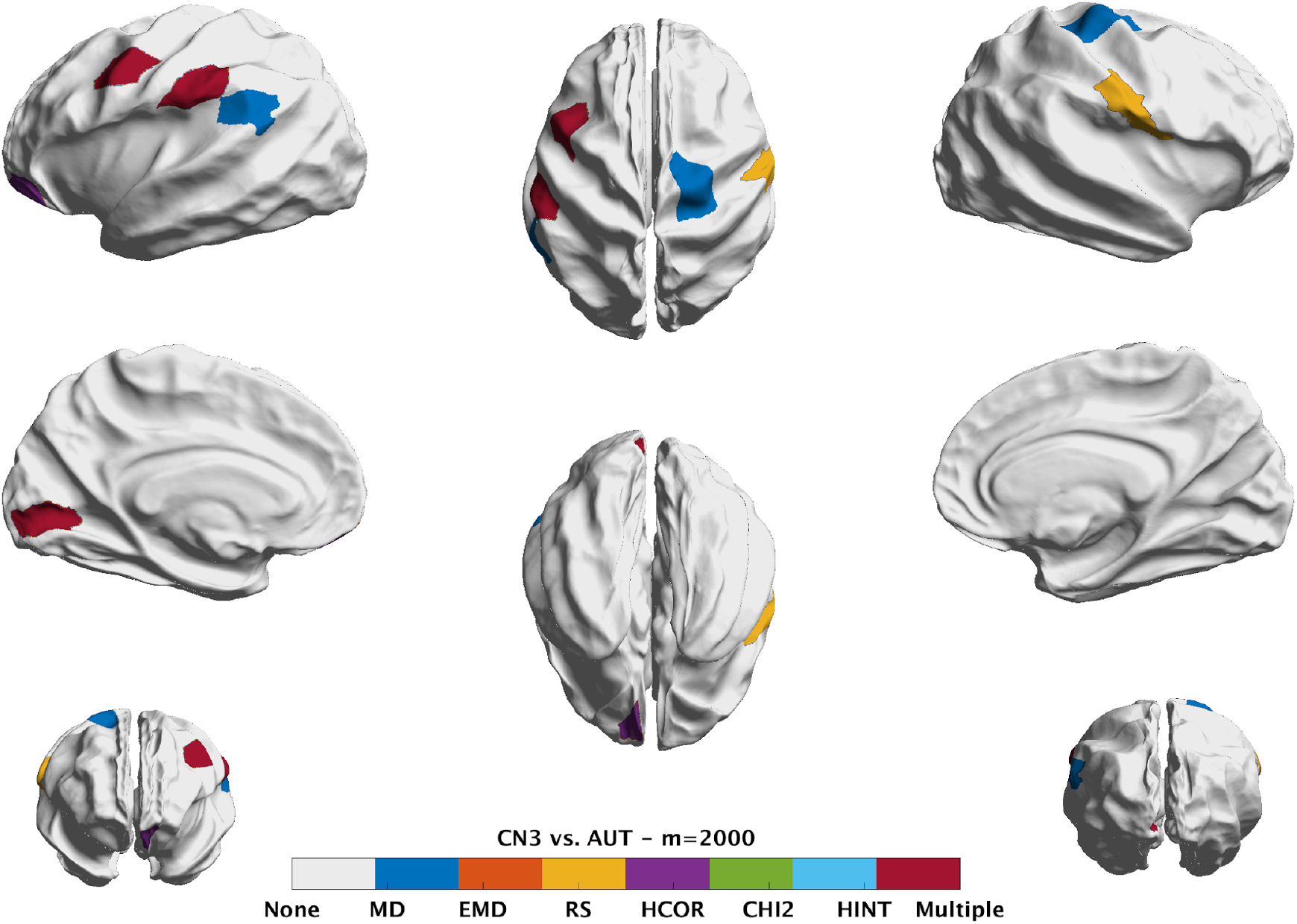
Visualization of the most discriminative regions as derived from the CN3 vs. AUT experiment at m=2000. These regions cover the lingual, supra-marginal, post- and precentral areas.

### Replication in AIBL dataset

In order to test whether the results and insights from this study on ADNI would generalize to a similar dataset, we’ve analyzed the AIBL dataset (see Table 2). The predictive utility for different combinations of edge weights and spatial scale (*m*) are shown in Figure 8. Although there are numerical differences in performance among different methods at fixed scale (*m*), their pattern remains similar across different spatial scales within the same dataset, as was observed in Fig 3 for the ADNI and ABIDE datasets. Based on posthoc statistical analyses (in the same fashion described earlier), we learn they are indeed not statistically significantly different from each other (the critical difference figure is not shown here to save space, as it is a single line connecting them all). That lack of significant differences is also true either for a fixed *m* (across different EW), or for a fixed EW (across different m).

**Fig. 8.**
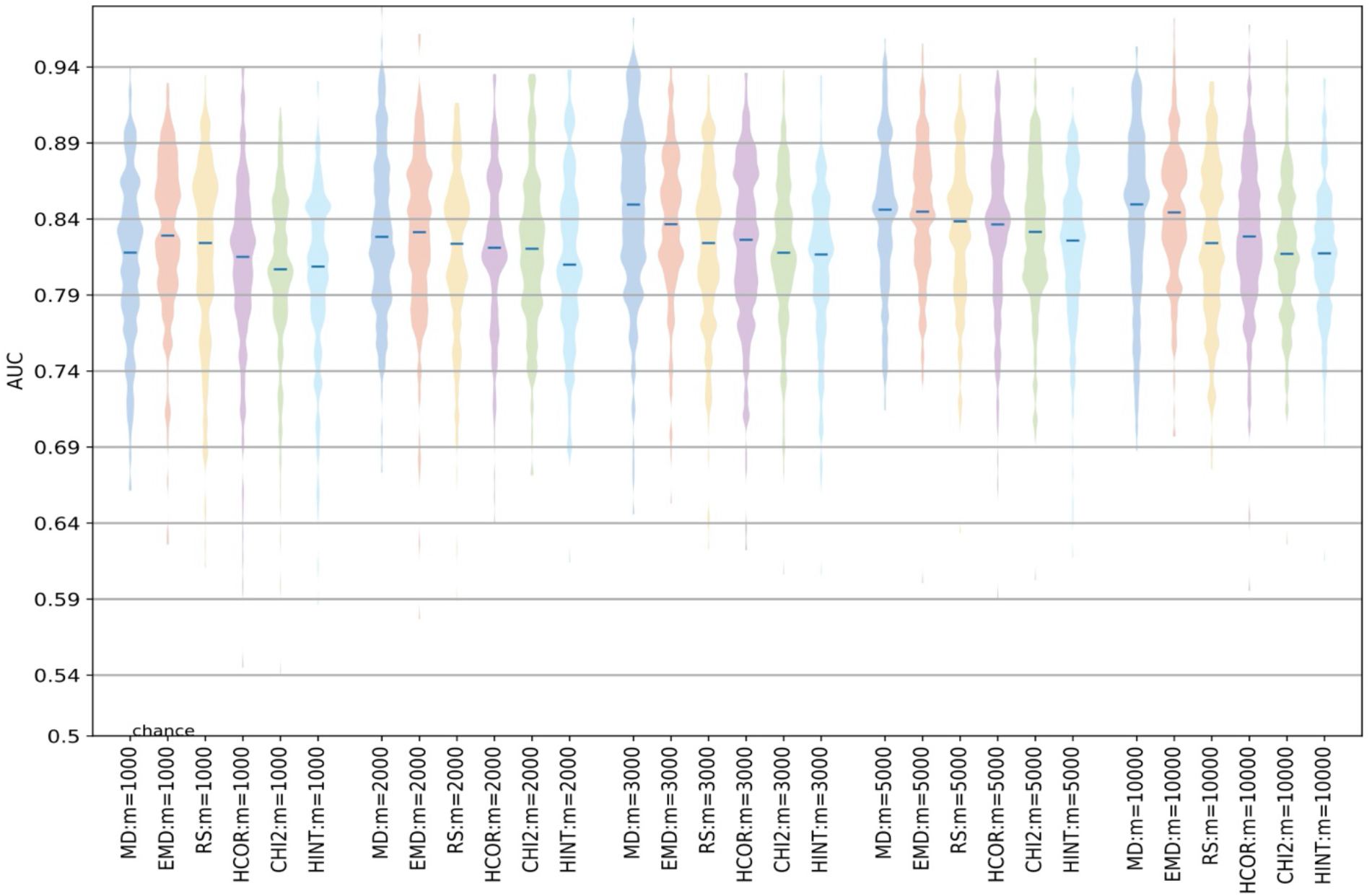
Classification performance for the different network methods (different edge weight metrics at different spatial resolutions of m) in discriminating AD2 from their respective control groups (CN4) from the AIBL dataset (see Table 2). The performance presented here is a distribution of AUC values from 200 randomized train/test splits of RHsT. It is clear that AUC for different combinations is quite similar to each other (as in Fig 3 for ADNI and ABIDE datasets). That is also true either for a fixed m (and different edge weights) or a fixed edge weight (and different values of m).

We note that the median AUC (to discriminate AD2 from CN4) across all combinations for the AIBL dataset is 0.83. This is in the typical range of NC vs. AD performance we notice in the Alzheimer’s literature, although lower than that noticed in the ADNI1 dataset of 0.87. This slight difference could be attributed to a number of factors, including a slightly different population, different feature extraction library (graynet relies on fully python-based stack), different machine learning library (scikit-learn based on libsvm vs. Matlab’s built-in SVM implementation), and most of all to a much smaller sample (n=131, which is only a third of the corresponding ADNI1 subset with n=412). That said, the patterns in performance observed in Fig. 8 i.e. lack of significant differences in performance for a fixed spatial scale (*m*), or for a fixed EW, replicate the main patterns from the ADNI1 dataset in the AIBL dataset.

### Future directions

While we present the results from a large number of experiments (n=90, 6 edge weights at 5 different scales *m* for the 3 datasets) covering two large publicly available datasets, two disease and age groups and three different levels of separability, there is certainly room for further analysis. Future studies could consider additional histogram distances, and performing the comparison with different types of classifiers (other than SVM such as linear discriminant or random forests).

It is possible that lack of sufficiently large sample size could be a contributor to the observed lack of statistically significant differences. This might especially be the case in challenging classification experiments such as CN3 vs. AUT. Moreover, given the multi-site nature of these large public datasets, properly accounting for the site and other relevant confounds would be worthy of further investigation. Such a broadening of scope for the study is not only computationally very intensive, but we believe studying the above is unlikely to change the conclusions. It would be nevertheless useful to quantitatively support it.

It would also be interesting to study the impact of different atlas choices (other than *fsaverage,* such as MNI152), parcellation (such as (Destrieux et al. 2010)) and subdivision schemes (functional or geometric or multimodal) (Eickhoff et al. 2015; Glasser et al. 2016), potential neuroimaging artifacts and confounds (Churchill et al. 2015; J. P. Lerch, van der Kouwe, and Raznahan 2017), but this would be demanding not only computationally but also in expert manpower for quality control (typically unavailable). It would also be quite interesting to replicate this study in the context of differential diagnosis (Raamana, Rosen, et al. 2014). A cross-modal comparison (Reid et al. 2015), in terms of predictive performance, with network-level features derived from modalities such as task-free fMRI would also be interesting.

## Conclusions

We have studied six different ways of constructing weighted networks derived from cortical thickness features, based on a novel method to derive edge weights based on histogram distances. We performed a comprehensive model comparison based on extensive cross-validation of their predictive utility and nonparametric statistical tests. This has been studied under three separabilities (ranging from pronounced, mild to subtle differences) derived from three independent and large publicly available datasets.

Some interesting results of this study based on the single-subject classification results are:

- the simpler methods of edge weight computation such as the difference in median thickness are as predictive as the sophisticated methods relying on the richer descriptions based on complete histograms.
- within a given method, the impact of a spatial scale *m* on predictive performance is not significant. The most popular way of computing edge weights in group-wise analysis i.e. histogram correlation, is shown to be the least predictive of disease-status in the context of individualized prediction via HiWeNet.

We have also developed and shared multiple open source toolboxes called graynet, visualqc and neuropredict to enable easy reuse of the methods and best practices presented in this study.

## Compliance with Ethical Standards

## Research involving Human Participants and/or Animals

Not applicable.

## Informed consent

Not applicable.

## ADNI

Data collection and sharing for this project was funded by the Alzheimer’s Disease Neuroimaging Initiative (ADNI) (National Institutes of Health Grant U01 AG024904) and DOD ADNI (Department of Defense award number W81XWH-12-2-0012). ADNI is funded by the National Institute on Aging, the National Institute of Biomedical Imaging and Bioengineering, and through generous contributions from the following: AbbVie, Alzheimer’s Association; Alzheimer’s Drug Discovery Foundation; Araclon Biotech; BioClinica, Inc.; Biogen; Bristol-Myers Squibb Company; CereSpir, Inc.; Cogstate; Eisai Inc.; Elan Pharmaceuticals, Inc.; Eli Lilly and Company; EuroImmun; F. Hoffmann-La Roche Ltd and its affiliated company Genentech, Inc.; Fujirebio; GE Healthcare; IXICO Ltd.; Janssen Alzheimer Immunotherapy Research & Development, LLC.; Johnson & Johnson Pharmaceutical Research & Development LLC.; Lumosity; Lundbeck; Merck & Co., Inc.; Meso Scale Diagnostics, LLC.; NeuroRx Research; Neurotrack Technologies; Novartis Pharmaceuticals Corporation; Pfizer Inc.; Piramal Imaging; Servier; Takeda Pharmaceutical Company; and Transition Therapeutics. The Canadian Institutes of Health Research is providing funds to support ADNI clinical sites in Canada. Private sector contributions are facilitated by the Foundation for the National Institutes of Health (www.fnih.org). The grantee organization is the Northern California Institute for Research and Education, and the study is coordinated by the Alzheimer’s Therapeutic Research Institute at the University of Southern California. ADNI data are disseminated by the Laboratory for NeuroImaging at the University of Southern California.

## AIBL

Data used in the preparation of this article was obtained from the Australian Imaging Biomarkers and Lifestyle flagship study of ageing (AIBL) funded by the Commonwealth Scientific and Industrial Research Organisation (CSIRO) which was made available at the ADNI database (www.loni.usc.edu/ADNI). The AIBL researchers contributed data but did not participate in analysis or writing of this report. AIBL researchers are listed at www.aibl.csiro.au.

## ABIDE I

Primary support for the work by Adriana Di Martino was provided by the NIMH (K23MH087770) and the Leon Levy Foundation. Primary support for the work by Michael P. Milham and the INDI team was provided by gifts from Joseph P. Healy and the Stavros Niarchos Foundation to the Child Mind Institute, as well as by an NIMH award to MPM (R03MH096321).

## Supplementary material

## Appendix A - Details of subjects used in this study

## Subject IDs from ADNI in Table 1

Note these subjects are all from baseline.

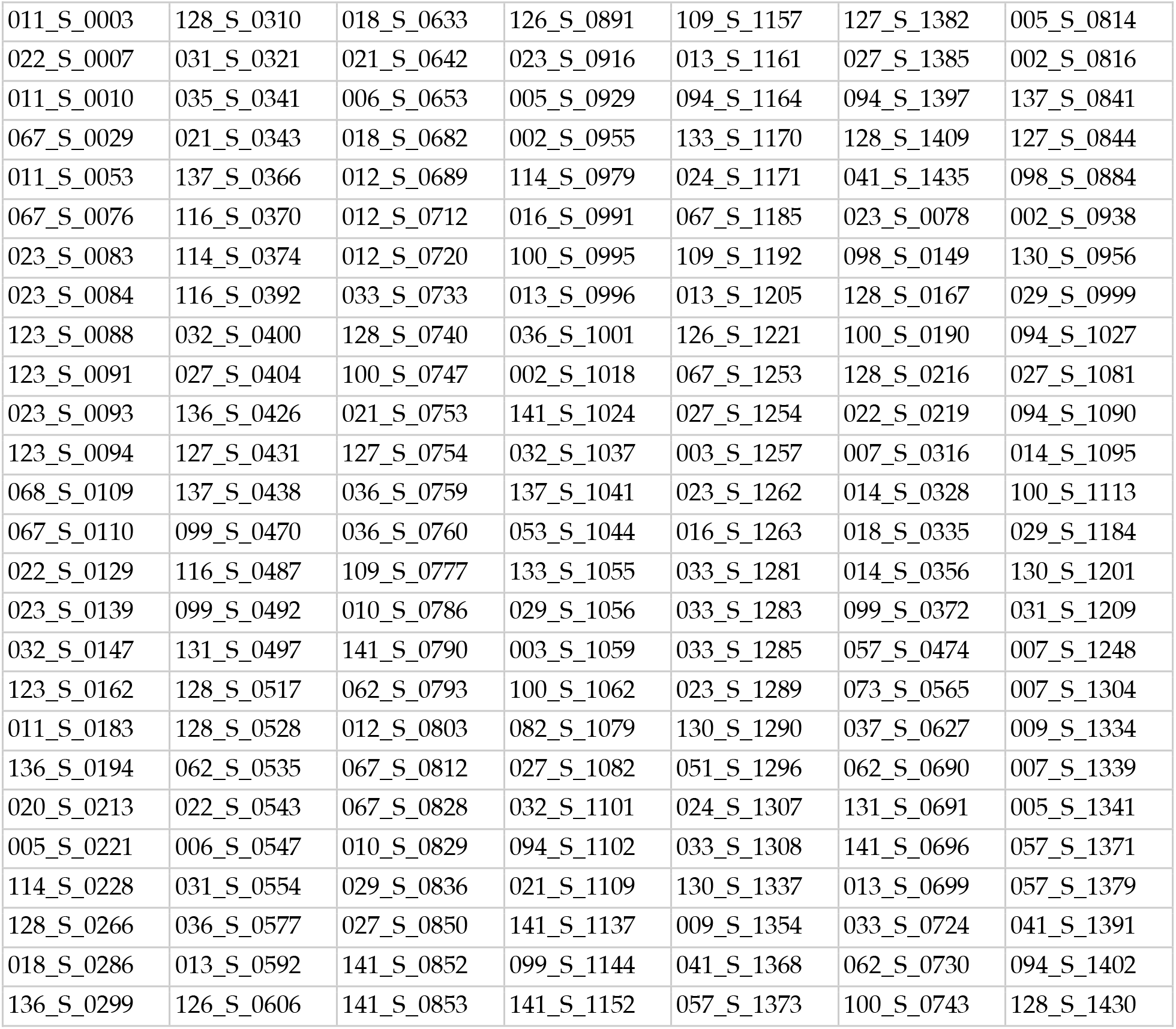
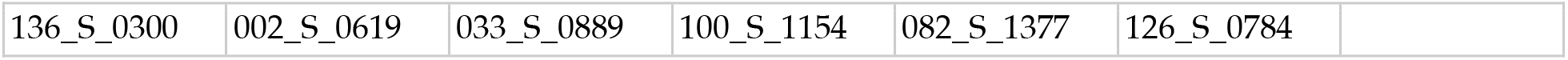
AD from Table 1.

AD from Table 1
|  |  |  |  |  |  |  |
| --- | --- | --- | --- | --- | --- | --- |
| 011_S_0003 | 128_S_0310 | 018_S_0633 | 126_S_0891 | 109_S_1157 | 127_S_1382 | 005_S_0814 |
| 022_S_0007 | 031_S_0321 | 021_S_0642 | 023_S_0916 | 013_S_1161 | 027_S_1385 | 002_S_0816 |
| 011_S_0010 | 035_S_0341 | 006_S_0653 | 005_S_0929 | 094_S_1164 | 094_S_1397 | 137_S_0841 |
| 067_S_0029 | 021_S_0343 | 018_S_0682 | 002_S_0955 | 133_S_1170 | 128_S_1409 | 127_S_0844 |
| 011_S_0053 | 137_S_0366 | 012_S_0689 | 114_S_0979 | 024_S_1171 | 041_S_1435 | 098_S_0884 |
| 067_S_0076 | 116_S_0370 | 012_S_0712 | 016_S_0991 | 067_S_1185 | 023_S_0078 | 002_S_0938 |
| 023_S_0083 | 114_S_0374 | 012_S_0720 | 100_S_0995 | 109_S_1192 | 098_S_0149 | 130_S_0956 |
| 023_S_0084 | 116_S_0392 | 033_S_0733 | 013_S_0996 | 013_S_1205 | 128_S_0167 | 029_S_0999 |
| 123_S_0088 | 032_S_0400 | 128_S_0740 | 036_S_1001 | 126_S_1221 | 100_S_0190 | 094_S_1027 |
| 123_S_0091 | 027_S_0404 | 100_S_0747 | 002_S_1018 | 067_S_1253 | 128_S_0216 | 027_S_1081 |
| 023_S_0093 | 136_S_0426 | 021_S_0753 | 141_S_1024 | 027_S_1254 | 022_S_0219 | 094_S_1090 |
| 123_S_0094 | 127_S_0431 | 127_S_0754 | 032_S_1037 | 003_S_1257 | 007_S_0316 | 014_S_1095 |
| 068_S_0109 | 137_S_0438 | 036_S_0759 | 137_S_1041 | 023_S_1262 | 014_S_0328 | 100_S_1113 |
| 067_S_0110 | 099_S_0470 | 036_S_0760 | 053_S_1044 | 016_S_1263 | 018_S_0335 | 029_S_1184 |
| 022_S_0129 | 116_S_0487 | 109_S_0777 | 133_S_1055 | 033_S_1281 | 014_S_0356 | 130_S_1201 |
| 023_S_0139 | 099_S_0492 | 010_S_0786 | 029_S_1056 | 033_S_1283 | 099_S_0372 | 031_S_1209 |
| 032_S_0147 | 131_S_0497 | 141_S_0790 | 003_S_1059 | 033_S_1285 | 057_S_0474 | 007_S_1248 |
| 123_S_0162 | 128_S_0517 | 062_S_0793 | 100_S_1062 | 023_S_1289 | 073_S_0565 | 007_S_1304 |
| 011_S_0183 | 128_S_0528 | 012_S_0803 | 082_S_1079 | 130_S_1290 | 037_S_0627 | 009_S_1334 |
| 136_S_0194 | 062_S_0535 | 067_S_0812 | 027_S_1082 | 051_S_1296 | 062_S_0690 | 007_S_1339 |
| 020_S_0213 | 022_S_0543 | 067_S_0828 | 032_S_1101 | 024_S_1307 | 131_S_0691 | 005_S_1341 |
| 005_S_0221 | 006_S_0547 | 010_S_0829 | 094_S_1102 | 033_S_1308 | 141_S_0696 | 057_S_1371 |
| 114_S_0228 | 031_S_0554 | 029_S_0836 | 021_S_1109 | 130_S_1337 | 013_S_0699 | 057_S_1379 |
| 128_S_0266 | 036_S_0577 | 027_S_0850 | 141_S_1137 | 009_S_1354 | 033_S_0724 | 041_S_1391 |
| 018_S_0286 | 013_S_0592 | 141_S_0852 | 099_S_1144 | 041_S_1368 | 062_S_0730 | 094_S_1402 |
| 136_S_0299 | 126_S_0606 | 141_S_0853 | 141_S_1152 | 057_S_1373 | 100_S_0743 | 128_S_1430 |

|  |  |  |  |  |  |
| --- | --- | --- | --- | --- | --- |
| 136_S_0300 | 002_S_0619 | 033_S_0889 | 100_S_1154 | 082_S_1377 | 126_S_0784 |

CN1 subjects used in Table 1.

CN1 subjects used in Table 1
|  |  |  |  |  |  |  |
| --- | --- | --- | --- | --- | --- | --- |
| 011_S_0002 | 123_S_0106 | 073_S_0312 | 013_S_0502 | 127_S_0684 | 003_S_0931 | 116_S_1249 |
| 011_S_0005 | 123_S_0113 | 072_S_0315 | 126_S_0506 | 002_S_0685 | 057_S_0934 | 052_S_1250 |
| 011_S_0008 | 027_S_0120 | 131_S_0319 | 033_S_0516 | 137_S_0686 | 052_S_0951 | 052_S_1251 |
| 022_S_0014 | 131_S_0123 | 037_S_0327 | 014_S_0519 | 094_S_0692 | 023_S_0963 | 082_S_1256 |
| 100_S_0015 | 041_S_0125 | 021_S_0337 | 128_S_0522 | 141_S_0717 | 109_S_0967 | 002_S_1261 |
| 011_S_0016 | 068_S_0127 | 099_S_0352 | 133_S_0525 | 141_S_0726 | 130_S_0969 | 094_S_1267 |
| 067_S_0019 | 035_S_0156 | 016_S_0359 | 094_S_0526 | 006_S_0731 | 137_S_0972 | 013_S_1276 |
| 011_S_0021 | 021_S_0159 | 116_S_0360 | 099_S_0534 | 033_S_0734 | 041_S_1002 | 002_S_1280 |
| 011_S_0022 | 114_S_0166 | 082_S_0363 | 016_S_0538 | 033_S_0741 | 012_S_1009 | 100_S_1286 |
| 011_S_0023 | 098_S_0171 | 018_S_0369 | 128_S_0545 | 009_S_0751 | 109_S_1013 | 020_S_1288 |
| 023_S_0031 | 098_S_0172 | 116_S_0382 | 014_S_0548 | 082_S_0761 | 109_S_1014 | 131_S_1301 |
| 100_S_0035 | 114_S_0173 | 073_S_0386 | 035_S_0555 | 141_S_0767 | 033_S_1016 | 035_S_0048 |
| 099_S_0040 | 067_S_0177 | 027_S_0403 | 014_S_0558 | 062_S_0768 | 036_S_1023 | 022_S_0066 |
| 018_S_0043 | 136_S_0184 | 126_S_0405 | 002_S_0559 | 057_S_0779 | 024_S_1063 | 022_S_0130 |
| 100_S_0047 | 136_S_0186 | 002_S_0413 | 013_S_0575 | 141_S_0810 | 033_S_1086 | 136_S_0196 |
| 067_S_0056 | 068_S_0210 | 114_S_0416 | 036_S_0576 | 036_S_0813 | 141_S_1094 | 073_S_0311 |
| 023_S_0058 | 005_S_0223 | 010_S_0419 | 062_S_0578 | 057_S_0818 | 033_S_1098 | 131_S_0441 |
| 067_S_0059 | 128_S_0229 | 010_S_0420 | 114_S_0601 | 009_S_0842 | 051_S_1123 | 014_S_0520 |
| 023_S_0061 | 128_S_0230 | 018_S_0425 | 005_S_0602 | 029_S_0843 | 012_S_1133 | 005_S_0553 |
| 010_S_0067 | 130_S_0232 | 133_S_0433 | 126_S_0605 | 029_S_0845 | 032_S_1169 | 116_S_0648 |
| 007_S_0068 | 128_S_0245 | 131_S_0436 | 005_S_0610 | 009_S_0862 | 023_S_1190 | 094_S_0711 |
| 100_S_0069 | 067_S_0257 | 037_S_0454 | 031_S_0618 | 128_S_0863 | 068_S_1191 | 129_S_0778 |
| 007_S_0070 | 127_S_0259 | 137_S_0459 | 127_S_0622 | 029_S_0866 | 941_S_1194 | 029_S_0824 |
| 123_S_0072 | 127_S_0260 | 037_S_0467 | 012_S_0637 | 109_S_0876 | 941_S_1195 | 003_S_0907 |
| 027_S_0074 | 041_S_0262 | 010_S_0472 | 082_S_0640 | 020_S_0883 | 941_S_1197 | 003_S_0981 |
| 023_S_0081 | 128_S_0272 | 032_S_0479 | 057_S_0643 | 130_S_0886 | 941_S_1202 | 021_S_0984 |
| 136_S_0086 | 137_S_0283 | 006_S_0484 | 021_S_0647 | 098_S_0896 | 941_S_1203 | 024_S_0985 |
| 073_S_0089 | 002_S_0295 | 133_S_0488 | 116_S_0657 | 041_S_0898 | 007_S_1206 | 003_S_1021 |
| 099_S_0090 | 123_S_0298 | 094_S_0489 | 036_S_0672 | 020_S_0899 | 007_S_1222 | 062_S_1099 |
| 032_S_0095 | 137_S_0301 | 133_S_0493 | 032_S_0677 | 033_S_0920 | 116_S_1232 | 130_S_1200 |
| 022_S_0096 | 037_S_0303 | 006_S_0498 | 126_S_0680 | 033_S_0923 | 094_S_1241 | 012_S_1212 |
| 020_S_0097 | 082_S_0304 | 128_S_0500 | 006_S_0681 | 023_S_0926 | 128_S_1242 | 023_S_1306 |

MCIc subjects used in Table 1.

MC1c subjects used in Table 1
|  |  |  |  |  |  |  |
| --- | --- | --- | --- | --- | --- | --- |
| 002_S_0954 | 023_S_0042 | 035_S_0204 | 127_S_0394 | 011_S_0861 | 041_S_0549 | 126_S_1077 |
| 002_S_1070 | 023_S_0388 | 035_S_0997 | 133_S_0638 | 011_S_1282 | 041_S_1412 | 127_S_1427 |
| 005_S_0222 | 023_S_0604 | 051_S_1331 | 136_S_0195 | 013_S_0325 | 041_S_1423 | 128_S_0947 |
| 007_S_0041 | 023_S_0855 | 052_S_0952 | 141_S_0982 | 014_S_0658 | 057_S_0941 | 130_S_0423 |
| 007_S_0128 | 023_S_0887 | 052_S_1054 | 941_S_1311 | 023_S_0030 | 057_S_1217 | 133_S_0727 |
| 007_S_0344 | 023_S_1247 | 053_S_0507 | 941_S_1363 | 023_S_0625 | 067_S_0045 | 133_S_0913 |
| 011_S_0856 | 027_S_0461 | 062_S_1299 | 002_S_0729 | 027_S_0179 | 067_S_0077 | 136_S_0695 |
| 013_S_0240 | 033_S_0723 | 067_S_0243 | 005_S_0572 | 027_S_0256 | 094_S_0434 | 141_S_0915 |
| 013_S_0860 | 033_S_0725 | 067_S_0336 | 006_S_1130 | 027_S_1213 | 098_S_0269 | 141_S_1244 |
| 022_S_0750 | 033_S_0906 | 094_S_1015 | 007_S_0249 | 027_S_1387 | 099_S_0054 | 941_S_1295 |
| 022_S_1394 | 033_S_0922 | 094_S_1398 | 011_S_0241 | 033_S_0567 | 099_S_0111 |  |

CN2 subjects used in Table 1.

CN2 subjects used in Table 1
|  |  |  |  |  |  |  |
| --- | --- | --- | --- | --- | --- | --- |
| 011_S_0002 | 020_S_0097 | 073_S_0386 | 002_S_0559 | 062_S_0768 | 041_S_1002 | 013_S_1276 |
| 011_S_0005 | 027_S_0120 | 027_S_0403 | 013_S_0575 | 057_S_0779 | 109_S_1013 | 002_S_1280 |
| 011_S_0008 | 131_S_0123 | 126_S_0405 | 036_S_0576 | 141_S_0810 | 109_S_1014 | 020_S_1288 |
| 022_S_0014 | 041_S_0125 | 002_S_0413 | 062_S_0578 | 036_S_0813 | 033_S_1016 | 131_S_1301 |
| 011_S_0016 | 035_S_0156 | 114_S_0416 | 114_S_0601 | 029_S_0843 | 036_S_1023 | 035_S_0048 |
| 067_S_0019 | 114_S_0166 | 133_S_0433 | 005_S_0602 | 029_S_0845 | 024_S_1063 | 022_S_0066 |
| 011_S_0021 | 098_S_0171 | 131_S_0436 | 126_S_0605 | 128_S_0863 | 033_S_1086 | 022_S_0130 |
| 011_S_0022 | 098_S_0172 | 006_S_0484 | 005_S_0610 | 029_S_0866 | 141_S_1094 | 136_S_0196 |
| 011_S_0023 | 114_S_0173 | 133_S_0488 | 127_S_0622 | 109_S_0876 | 033_S_1098 | 073_S_0311 |
| 023_S_0031 | 067_S_0177 | 094_S_0489 | 082_S_0640 | 020_S_0883 | 051_S_1123 | 131_S_0441 |
| 099_S_0040 | 136_S_0184 | 133_S_0493 | 057_S_0643 | 130_S_0886 | 023_S_1190 | 014_S_0520 |
| 067_S_0056 | 136_S_0186 | 006_S_0498 | 036_S_0672 | 098_S_0896 | 941_S_1194 | 005_S_0553 |
| 023_S_0058 | 130_S_0232 | 013_S_0502 | 126_S_0680 | 041_S_0898 | 941_S_1195 | 094_S_0711 |
| 067_S_0059 | 067_S_0257 | 126_S_0506 | 006_S_0681 | 020_S_0899 | 941_S_1197 | 029_S_0824 |
| 023_S_0061 | 127_S_0259 | 033_S_0516 | 127_S_0684 | 033_S_0920 | 941_S_1202 | 003_S_0907 |
| 007_S_0068 | 127_S_0260 | 014_S_0519 | 002_S_0685 | 033_S_0923 | 941_S_1203 | 003_S_0981 |
| 007_S_0070 | 002_S_0295 | 133_S_0525 | 094_S_0692 | 023_S_0926 | 007_S_1206 | 024_S_0985 |
| 027_S_0074 | 082_S_0304 | 094_S_0526 | 141_S_0717 | 003_S_0931 | 007_S_1222 | 003_S_1021 |
| 023_S_0081 | 073_S_0312 | 099_S_0534 | 006_S_0731 | 057_S_0934 | 052_S_1250 | 062_S_1099 |
| 136_S_0086 | 131_S_0319 | 016_S_0538 | 033_S_0734 | 052_S_0951 | 052_S_1251 | 130_S_1200 |
| 073_S_0089 | 099_S_0352 | 014_S_0548 | 033_S_0741 | 023_S_0963 | 082_S_1256 | 023_S_1306 |
| 099_S_0090 | 016_S_0359 | 035_S_0555 | 082_S_0761 | 109_S_0967 | 002_S_1261 |  |
| 022_S_0096 | 082_S_0363 | 014_S_0558 | 141_S_0767 | 130_S_0969 | 094_S_1267 |  |

## Subject IDs excluded from the ADNI cohort

owing to failure in Freesurfer processing or other errors

|  |  |  |  |
| --- | --- | --- | --- |
| 006_S_0322 | 018_S_0277 | 062_S_1091 | 109_S_0840 |
| 006_S_0521 | 027_S_0948 | 067_S_0020 | 128_S_0701 |
| 010_S_0662 | 027_S_1335 | 067_S_0024 | 128_S_0805 |
| 010_S_0788 | 031_S_0773 | 073_S_1207 | 128_S_1181 |
| 011_S_0326 | 033_S_0888 | 094_S_0964 | 130_S_0460 |
| 014_S_0357 | 033_S_1087 | 100_S_0893 | 141_S_0340 |

## Subjects IDs used from ABIDE in this study

CN3 subjects used in Table 3.

**CN3 subjects used in Table 3**
|  |  |  |  |  |
| --- | --- | --- | --- | --- |
| Pitt_0050038 | Pitt_0050047 | UM_2_0050428 | UCLA_1_0051281 | KKI_0050776 |
| Pitt_0050039 | Pitt_0050046 | UM_2_0050426 | Caltech_0051484 | UM_1_0050334 |
| NYU_0051036 | Trinity_0051142 | UM_2_0050424 | SBL_0051562 | UM_1_0050335 |
| Pitt_0050034 | Leuven_1_0050703 | Leuven_2_0050730 | NYU_0051090 | Yale_0050577 |
| Pitt_0050035 | Leuven_1_0050701 | Leuven_2_0050731 | Leuven_1_0050699 | SBL_0051566 |
| Pitt_0050036 | NYU_0051105 | Leuven_2_0050732 | NYU_0051039 | NYU_0051041 |
| Pitt_0050037 | NYU_0051104 | Leuven_2_0050733 | UM_2_0050417 | Yale_0050558 |
| Pitt_0050030 | NYU_0051107 | Leuven_2_0050735 | Olin_0050122 | UM_1_0050366 |
| Pitt_0050031 | NYU_0051106 | Leuven_2_0050736 | Yale_0050553 | Trinity_0051137 |
| Pitt_0050032 | NYU_0051101 | Leuven_2_0050737 | Olin_0050109 | Pitt_0050050 |
| Pitt_0050033 | NYU_0051100 | Leuven_2_0050738 | SBL_0051561 | UCLA_2_0051309 |
| Pitt_0050049 | NYU_0051103 | Leuven_2_0050739 | Trinity_0051141 | UCLA_2_0051306 |
| Pitt_0050048 | NYU_0051102 | MaxMun_a_0051370 | UM_2_0050414 | UCLA_1_0051272 |
| NYU_0051038 | NYU_0051109 | Leuven_2_0050741 | Leuven_2_0050722 | NYU_0051079 |
| Pitt_0050041 | KKI_0050782 | Leuven_2_0050740 | UM_2_0050422 | USM_0050436 |
| Pitt_0050040 | USM_0050463 | Leuven_2_0050742 | UM_2_0050421 | USM_0050433 |
| Pitt_0050043 | USM_0050466 | Trinity_0050257 | UM_2_0050427 | Olin_0050119 |
| Pitt_0050042 | USM_0050467 | KKI_0050820 | UM_2_0050425 | Trinity_0050266 |
| Pitt_0050045 | USM_0050468 | SBL_0051567 | NYU_0051129 | SDSU_0050196 |
| Pitt_0050044 | USM_0050469 | SBL_0051564 | NYU_0051084 | Olin_0050113 |

AUT subjects used in Table 3.

AUT subjects used in Table 3
|  |  |  |  |  |
| --- | --- | --- | --- | --- |
| NYU_0051032 | UM_1_0050296 | UCLA_1_0051224 | MaxMun_d_0051350 | UM_1_0050326 |
| NYU_0051034 | Pitt_0050028 | UCLA_1_0051225 | NYU_0050985 | NYU_0050984 |
| UCLA_1_0051240 | Pitt_0050027 | UCLA_1_0051226 | USM_0050523 | Leuven_2_0050753 |
| Leuven_1_0050702 | NYU_0051028 | USM_0050488 | NYU_0050994 | Leuven_2_0050751 |
| USM_0050509 | Leuven_1_0050694 | USM_0050487 | NYU_0050997 | Leuven_2_0050757 |
| USM_0050505 | CMU_a_0050654 | Leuven_2_0050748 | NYU_0051008 | Leuven_2_0050754 |
| USM_0050501 | Leuven_1_0050711 | KKI_0050801 | NYU_0051009 | NYU_0050954 |
| USM_0050500 | USM_0050518 | KKI_0050802 | NYU_0051006 | NYU_0050956 |
| UM_1_0050315 | UM_1_0050278 | KKI_0050804 | NYU_0051007 | UCLA_2_0051302 |
| Leuven_2_0050746 | UM_1_0050308 | Pitt_0050011 | NYU_0051001 | CMU_b_0050652 |
| Trinity_0050251 | UM_2_0050410 | Pitt_0050016 | UCLA_1_0051237 | Pitt_0050007 |
| Trinity_0050250 | USM_0050531 | Pitt_0050015 | UCLA_1_0051235 | Pitt_0050003 |
| UCLA_1_0051219 | Trinity_0050249 | UCLA_1_0051231 | UCLA_1_0051234 | USM_0050532 |
| UCLA_1_0051218 | Trinity_0050242 | USM_0050520 | Pitt_0050057 | UCLA_1_0051209 |
| UCLA_1_0051215 | Trinity_0050245 | USM_0050525 | Pitt_0050055 | UCLA_1_0051206 |
| UCLA_1_0051214 | Trinity_0050246 | NYU_0050987 | Pitt_0050053 | UCLA_1_0051205 |
| USM_0050491 | SBL_0051585 | NYU_0050986 | NYU_0050998 | UCLA_1_0051201 |
| USM_0050493 | NYU_0051012 | UM_2_0050402 | Caltech_0051468 | NYU_0050989 |
| USM_0050492 | UCLA_2_0051317 | UM_2_0050406 | UM_1_0050321 | UM_1_0050285 |
| UM_1_0050298 | UCLA_1_0051223 | UCLA_2_0051293 | UM_1_0050320 | Leuven_1_0050689 |

## Subjects IDs used from the AIBL dataset

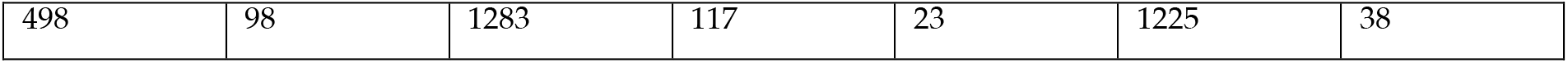
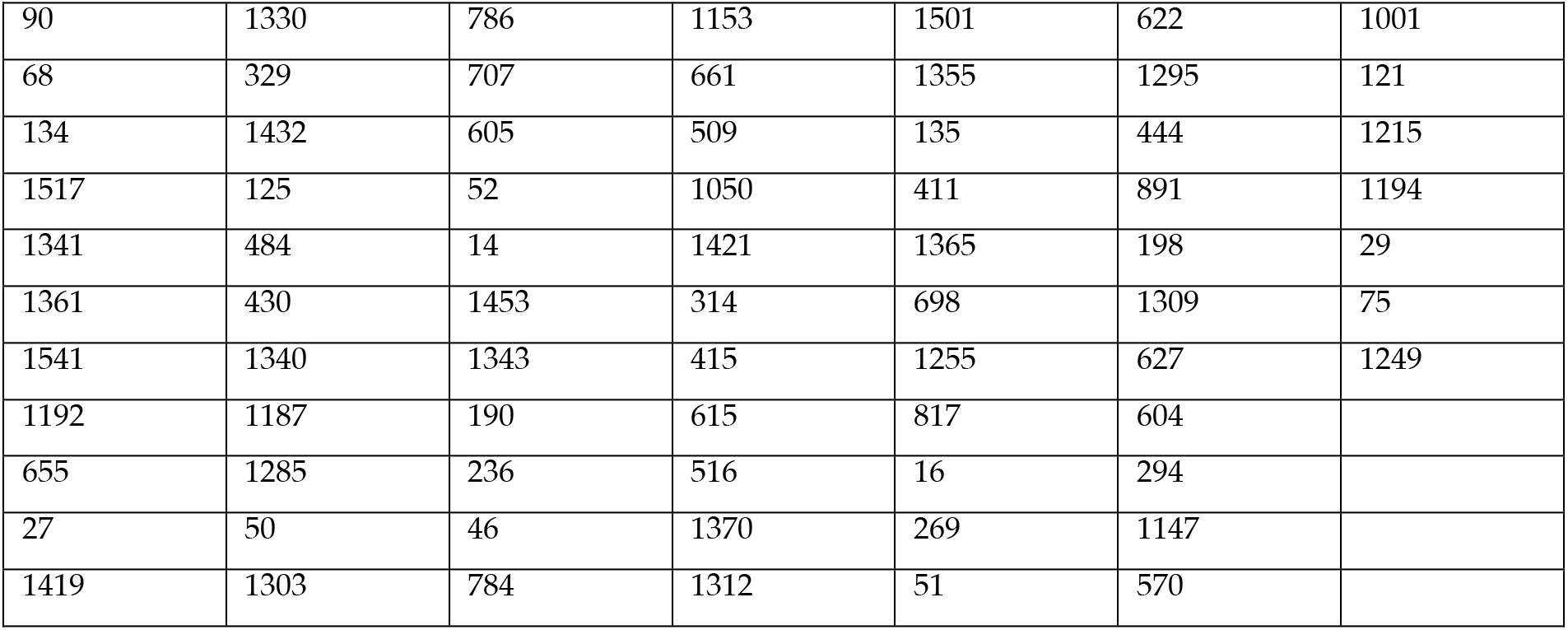
CN4: Baseline healthy controls.

CN4: Baseline healthy controls
|  |  |  |  |  |  |  |
| --- | --- | --- | --- | --- | --- | --- |
| 498 | 98 | 1283 | 117 | 23 | 1225 | 38 |
| 90 | 1330 | 786 | 1153 | 1501 | 622 | 1001 |
| 68 | 329 | 707 | 661 | 1355 | 1295 | 121 |
| 134 | 1432 | 605 | 509 | 135 | 444 | 1215 |
| 1517 | 125 | 52 | 1050 | 411 | 891 | 1194 |
| 1341 | 484 | 14 | 1421 | 1365 | 198 | 29 |
| 1361 | 430 | 1453 | 314 | 698 | 1309 | 75 |
| 1541 | 1340 | 1343 | 415 | 1255 | 627 | 1249 |
| 1192 | 1187 | 190 | 615 | 817 | 604 |  |
| 655 | 1285 | 236 | 516 | 16 | 294 |  |
| 27 | 50 | 46 | 1370 | 269 | 1147 |  |
| 1419 | 1303 | 784 | 1312 | 51 | 570 |  |

AD2: Baseline Alzheimer’s.

AD2: Baseline Alzheimer's
|  |  |  |  |  |  |  |  |
| --- | --- | --- | --- | --- | --- | --- | --- |
| 371 | 838 | 1135 | 1273 | 19 | 819 | 665 | 102 |
| 1316 | 567 | 1264 | 1013 | 1437 | 1549 | 1046 | 952 |
| 1484 | 1089 | 1482 | 1442 | 1547 | 1056 | 1513 |  |
| 488 | 893 | 1504 | 494 | 919 | 1032 | 978 |  |
| 1122 | 510 | 373 | 1457 | 361 | 970 | 1078 |  |
| 917 | 1559 | 417 | 1575 | 1090 | 1209 | 895 |  |
| 1577 | 1368 | 1260 | 993 | 1537 | 259 | 1066 |  |

## Appendix B

## The need for a trimmed estimator

In the Methods section, when describing the computation behind HiWeNet, we note that we remove 5% outliers from both tails of the thickness value distribution. The need to trim the distribution arises from the presence of several outlying values as can be seen from Fig. B1. There are a large number of vertices with zero and very small values (which are zoomed-in in the right panel in Figure B) as well as few unnaturally large values (over 6mm), making it necessary to trim the patch-wise distributions to stabilize the distance estimates between a random pair of patch-wise histograms. We observe similar trends across all patch sizes (all values of *m*).

**Figure B1:**
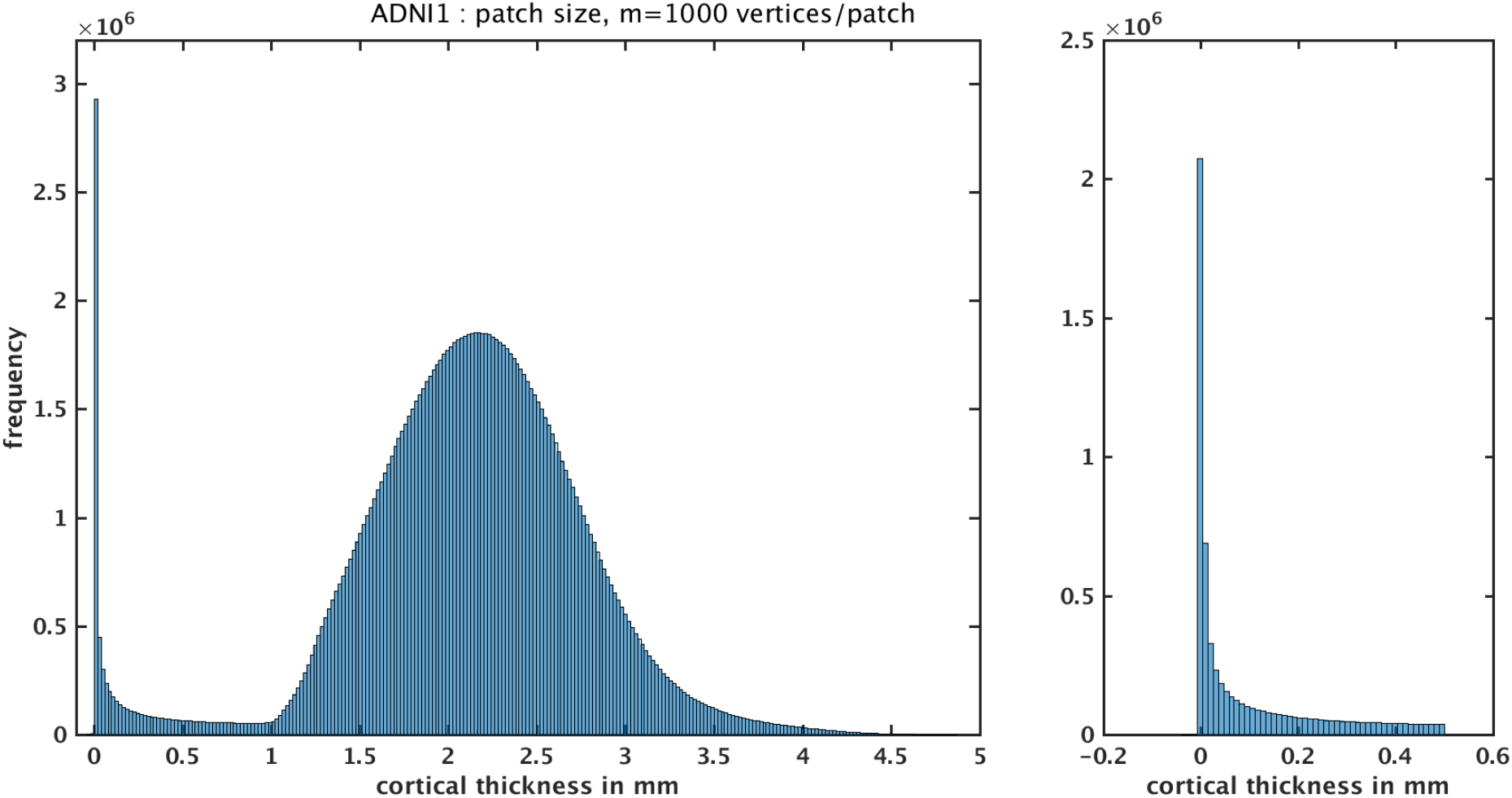
the full distribution of thickness values from ADNI1 dataset using all the subjects (CN1, CN2, MCI and AD) included in this study. It is clear there are a large number of vertices with zero and very small values, making it necessary to trim the patch-wise distributions to stabilize the distance estimates between a random pair of patch-wise histograms.

## Appendix C

To explore alternative representations for network-level features, we have extracted the following features: 1) compute a histogram for the distribution of thickness values from the entire cortex (‘grand histogram’), and 2) for each patch, represent its value by the histogram distance between its own histogram and the grand mean histogram. Let’s denote this method ‘relative_to_all’. This method results in a vector of length *n* only (number of patches for a given *m*) as opposed to fully-pairwise method adopted in this paper which results in *n*(n-1)/2* features. To understand their utility, we have evaluated their predictive performance for the 30 different feature sets based on ‘relative_to_all’ edge weight. Their performance did not differ substantially from the fully-pairwise network-level counterparts - see the figure below. The median baseline performance (median of the 30 median AUCs each from 200 CV repetitions) is at AUC=0.89 in the CN1 vs. AD experiment (compared to AUC=0.87 for the fully-pairwise network features), at AUC= 0.77 (compared to AUC=0.75) for CN2 vs. MCIc and at AUC=0.56 (compared to AUC=0.6) for CN3 vs. AUT. Although the simpler relative_to_all method seems to perform just as well or slightly numerically better when the differences are pronounced (CN1 vs. AD and MCIc), it does slightly worse in the more challenging experiment (CN3 vs. AUT). This is consistent with our previous experience wherein fully-pairwise network-level features performed increasingly better as the predictive challenge increased with decreasing separability (Raamana et al. 2015). These results are now included in Appendix C.

We’ve also updated our open source *hiwenet* package to provide this feature, crediting this reviewer for the idea (anonymously).

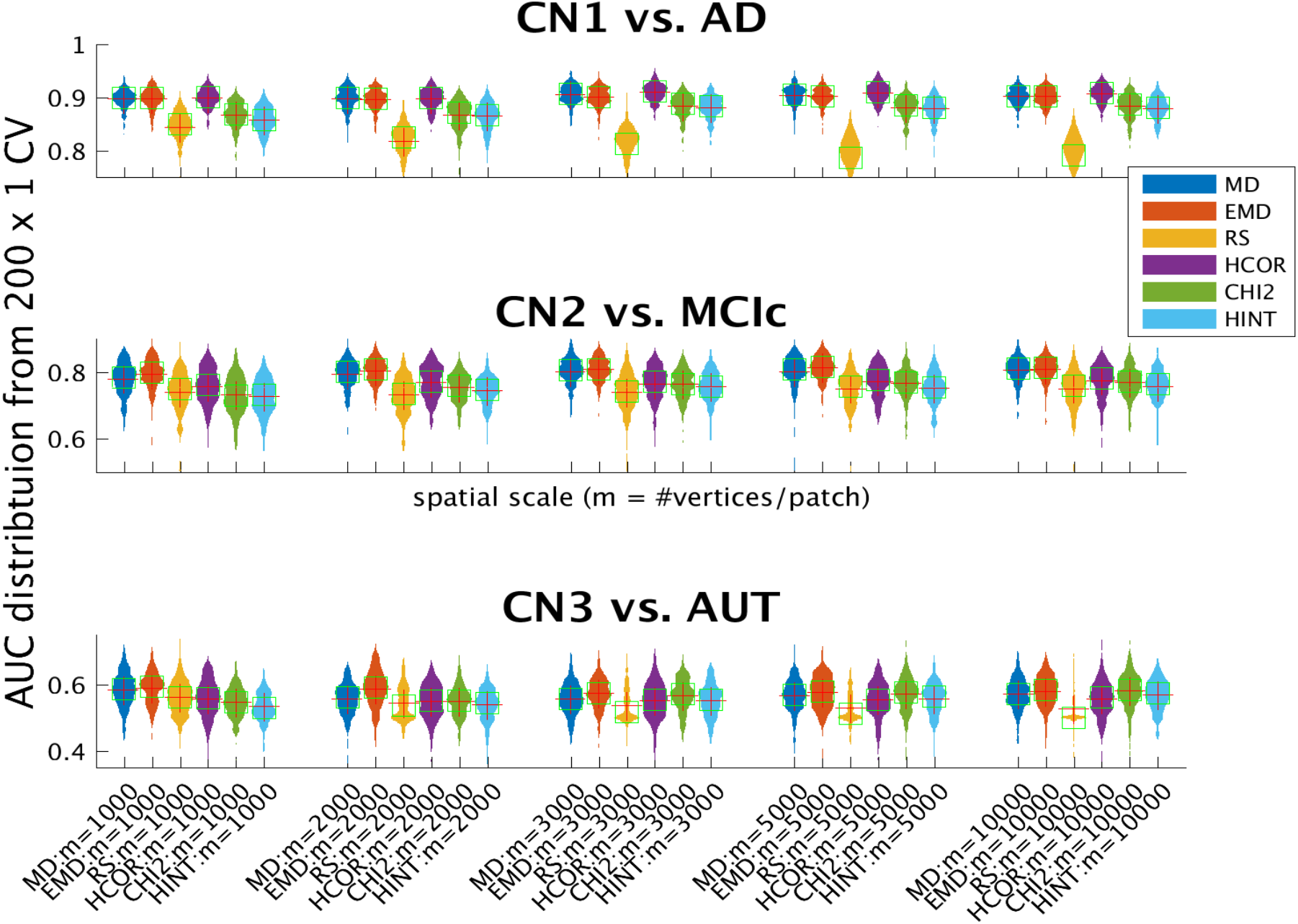

## Appendix D

The sites represented per diagnostic group in the ABIDE dataset are shown in the table below:

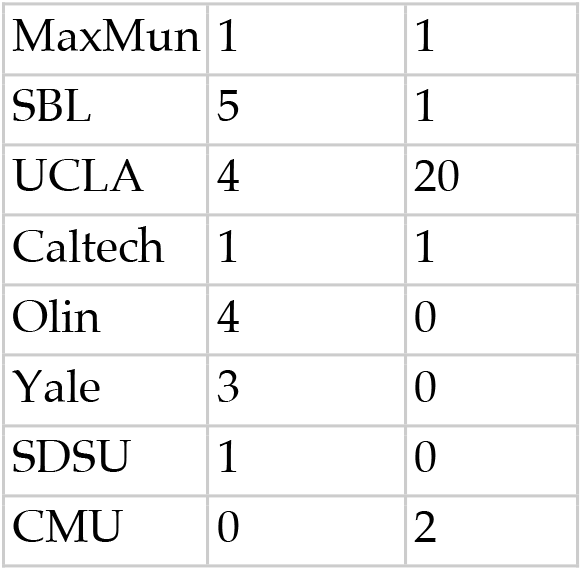

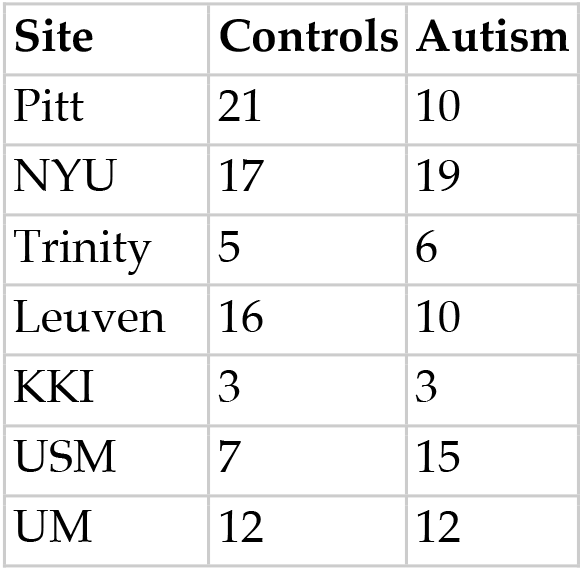

